# Adrenergic signaling in muscularis macrophages limits neuronal death following enteric infection

**DOI:** 10.1101/556340

**Authors:** Fanny Matheis, Paul A. Muller, Christina Graves, Ilana Gabanyi, Zachary J. Kerner, Diego Costa-Borges, Daniel Mucida

## Abstract

Enteric–associated neurons (EANs) are closely associated with immune cells and continuously monitor and modulate homeostatic intestinal functions, including motility. Bidirectional interactions between immune and neuronal cells are altered during disease processes such as neurodegeneration or irritable bowel syndrome. We investigated how infection-induced inflammation affects intrinsic EANs and the role of intestinal *muscularis* macrophages (MMs) in this process. Using murine model of bacterial infection, we observed long-term gastrointestinal symptoms including reduced motility and subtype-specific neuronal loss. Neuronal-specific translational–profiling uncovered a caspase 11–dependent EAN cell–death mechanism induced by enteric infections. MMs responded to luminal infection by upregulating a neuroprotective program; gain– and loss–of-function experiments indicated that β2-adrenergic receptor (β_2_-AR) signaling in MMs mediates neuronal protection during infection via an arginase 1-polyamine axis. Our results identify a mechanism of neuronal cell death post–infection and point to a role for tissue–resident MMs in limiting neuronal damage.

## Introduction

The gastrointestinal (GI) tract comprises the largest environmental interface of the body and is posed with the unique challenge of maintaining tolerance to dietary and microbial antigens while remaining poised to protect against pathogen invasion. Coordinated resistance and tolerance mechanisms serve to prevent pathogenic dissemination, limit excessive GI damage, and initiate recovery responses induced by pathogenic burden or injury. Rampant infection-induced inflammation is linked to immunopathology in several disease conditions, including cerebral malaria and respiratory viral infections (Medzhitov et al., 2012; Soares et al., 2014). In the GI tract, enteric infections often result in functional gastrointestinal disorders post pathogen clearance, and often enteric neurons, which occupy the intestinal tissue in large numbers, are targeted (Balemans et al., 2017). Clinical observations indicate that between 6-17% of individuals with irritable bowel syndrome (IBS) develop symptoms after episodes of enteric infections; while around 10% of people with bacterial gastroenteritis develop IBS (Holschneider et al., 2011; Ohman and Simren, 2010). Further, flavivirus strains with neuronal tropism, including West Nile and Zika virus, have also been reported to induce long-term intestinal dysmotility (White et al., 2018). The clinical presentation of post-infectious IBS includes unresolved low-grade intestinal inflammation, gastrointestinal motility impairment, and nerve damage (Beatty et al., 2014; Holschneider et al., 2011). However, the underlying mechanisms involved in infection–induced neuronal damage are incompletely understood.

The intestinal immune and nervous systems sense and integrate luminal cues, and regulate physiological processes, including gastrointestinal motility. Recent evidence suggests that intestinal–resident macrophage populations play a role in normal functioning of enteric neurons in the absence of infections (De Schepper et al., 2018; Muller et al., 2014). Specifically, *muscularis* macrophages (MMs), located within and surrounding the myenteric plexus, were shown to regulate the activity of enteric neurons and peristalsis via secretion of BMP2 in a microbiota– dependent manner (Muller et al., 2014). Additionally, we previously reported that in the unperturbed state, MMs displayed a tissue-protective gene-expression profile, a signature that was enhanced following enteric infection, and involved β_2_ adrenergic receptor (β_2_-AR) signaling (Gabanyi et al., 2016). It remained to then be determined, however, whether this neuron-macrophage crosstalk plays a role in infection–induced neuronal damage, or prevention thereof.

By utilizing imaging, cell sorting–independent transcriptomics, and pharmacological and genetic gain– and loss–of-function approaches, we report that murine enteric infection results in a rapid and persistent loss of intrinsic enteric-associated neurons (iEANs), which is associated with long–term gastrointestinal changes including intestinal dysmotility. Confocal imaging strategies revealed a subtype–specific neuronal loss, and transcriptomics and genetic approaches uncovered an iEAN cell death mechanism that is dependent on components of the inflammasome pathway. Depletion of MMs, located in close proximity to enteric neurons, resulted in enhanced infection-induced neuronal loss, suggesting a functional role for a MM tissue protective program induced upon infection. Myeloid–specific targeting of β_2_-AR, as well as arginase 1 (Arg1), coupled with pharmacological rescue experiments implicated MM-adrenergic signaling in post-infectious enteric neuronal protection via the production of polyamines. Our data identify a functional role for neuron–macrophage interactions in limiting infection-induced neuronal damage.

## Results

### Enteric pathogens trigger long-term impairment in GI physiology

Acute bacterial infections, including *Salmonella enterica* serovar Typhimurium*, Shigella dysenteriae, and Campylobacter spp*. have previously been linked to long-term GI dysfunction resembling IBS (Ohman and Simren, 2010). To characterize functional outcomes of acute intracellular bacterial infection in mice, we used an attenuated strain of *Salmonella* Typhimurium, *spiB* (Tsolis et al, 1999), which harbors a mutation in the type-III secretion system, impacting its intracellular replication. We chose to use the *spiB* strain due to the fact that wild-type *Salmonella* Typhimurium rapidly invades and damages the intestinal wall, ultimately resulting in mortality in wild-type C57BL/6 mice, interfering with our goal of studying long-term functional intestinal changes post pathogen clearance. We observed that orally administered *spiB* was undetected in the feces by 7-10 days post infection (dpi) (Figure 1A, B). Infection with *spiB* caused mild intestinal inflammation as evidenced by increased levels of fecal lipocalin-2 (Lcn-2), that peaked at 6 dpi, just before *spiB* clearance, but remained elevated following pathogen clearance (Figure 1C). This persistent inflammatory response was associated with lasting functional GI changes, including increased ileal ring contractility and a persistent delay in gastrointestinal transit time (GITT) (Figure 1D, E).

**Figure 1:**
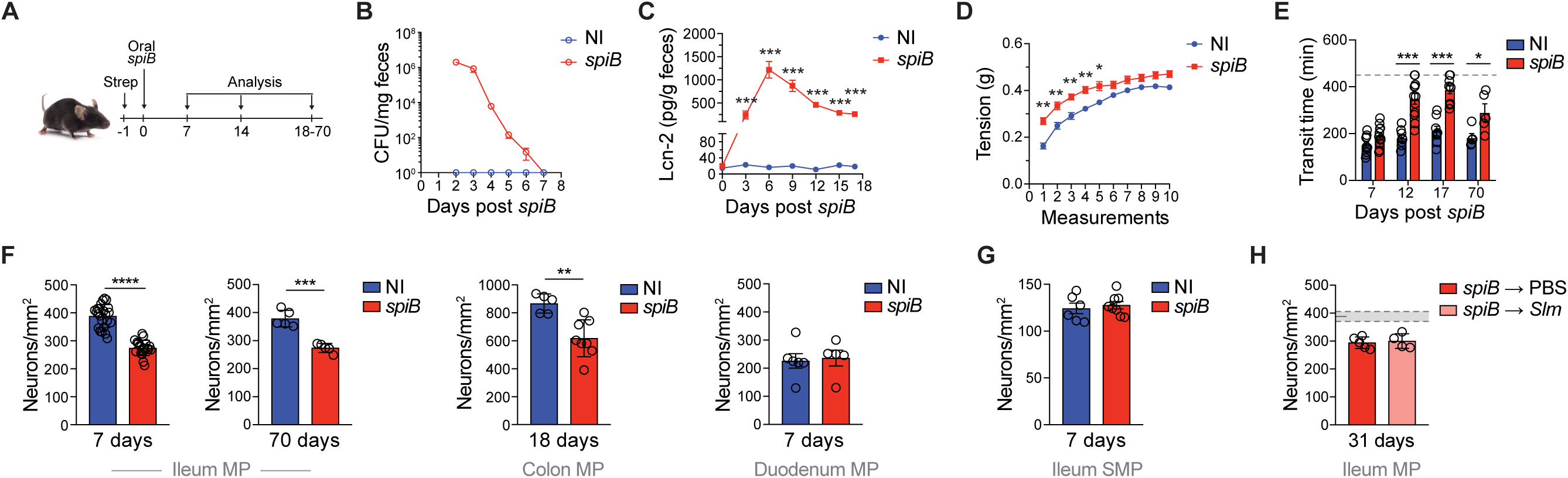
Inflammation induced by enteric infections triggers gut neuronal loss and dysmotility. (A) Experimental design for (B-J). (B-H) C57BL/6J mice were orally gavaged with PBS (non-infected, NI) or 10^9^ colony forming units (CFU) of *Salmonella spiB*. (B) Quantification of fecal CFU. (C) Quantification of fecal lipocalin-2 (Lcn-2) levels as assessed by ELISA post infection. (D) Ileal ring myography assessed on day 18 post infection. (E) Gastrointestinal motility as assessed by total gastrointestinal transit time evaluated on the indicated days post infection. Experiments were ended at 450 min (dashed line); (F-H) Neuronal quantification as assessed by immunofluorescence staining (ANNA-1) on the indicated days post infection in the (F) myenteric plexus of ileal segments (left), myenteric plexus of colonic segments (middle), myenteric plexus of duodenal segments (right); (H) submucosal plexus of ileal segments. (J) Neuronal quantification as assessed by immunofluorescence staining (ANNA-1) in the myenteric plexus of ileal segments of C57BL/6J mice orally gavaged PBS or 10^6^ CFU of wild-type *Salmonella* Typhimurium (Slm) 21 days post *spiB* infection, and sacrificed on day 10 post re-infection. Shaded area bounded by dashed lines indicates mean day 7 iEAN numbers +/- SEM of all control C57BL6/J mice. MP – Myenteric plexus; SMP – Submucosal plexus; Data are representative of at least 5 mice per condition. At least 2 independent experiments were performed. Data were analyzed by unpaired Student’s t-test or ANOVA with Bonferroni correction and are shown as average ± SEM; n.s. - not significant, *p ≤ 0.05, **p ≤ 0.01, ***p ≤ 0.001, ****p ≤ 0.0001

Changes in contractility and GITT can be associated with altered neuronal activity and damage (Travagli and Anselmi, 2016). Intrinsic enteric-associated neurons (iEANs), commonly referred to as the enteric nervous system, are neural crest–derived and organized in two distinct networks, the submucosal or Meissner’s plexus and myenteric or Auerbach’s plexus (Furness et al., 2013). In order to determine the impact of intestinal infection on iEANs, we quantified enteric neurons along the GI tract, in both the submucosal and myenteric plexuses. Following *spiB* infection, we observed a 20-30% reduction in myenteric neuron number as early as 7 dpi in the ileum and colon, both of which are major sites of *Salmonella* invasion; reduced neuronal counts were observed up to day 70 post–infection (Figure 1F). In contrast, neuronal numbers were preserved in the sites where *Salmonella* invasion normally does not occur, including the proximal small intestine (Figure 1F). We did not observe reduction in neuronal numbers in the submucosal plexus from mice infected with *spiB*; additionally, heat-killed *spiB* did not result in neuronal loss in the myenteric plexus (Figure 1G and Figure S1A). Loss of enteric neurons was also observed after infection with another bacterial pathogen, *Yersinia pseudotuberculosis*, previously reported to cause long-term restructuring of gut lymphatics known as immunological scaring (Fonseca et al., 2015) (Figure S1B). We further interrogated whether we could recapitulate our findings in the context of the clinically relevant protozoan pathogens *Toxoplasma gondii* (*T. gondii*) and *Trypanosoma cruzi* (*T. cruzi*), which are known to induce various GI-related pathologies (Dutra et al., 2009; Wilhelm and Yarovinsky, 2014). These protozoans also induced intestinal inflammation, increased ileal ring contractility, delayed GITT and significant neuronal loss (Figure S1C-G and *data not shown*). Taken together, these data indicate that iEAN loss may be a conserved feature of enteric infections, however we chose to continue with *spiB* as an infection model due to the inherent neurotropism of *T. gondii* and *T. cruzi*.

While it is generally accepted that enteric neurogenesis ceases in postnatal mammals, recent work suggests that Nestin+ stem cells may continuously replenish iEANs (Kulkarni et al., 2017). However, in the present study, we were unable to detect iEAN number recovery up to two months following *spiB* clearance (Figure 1F), nor did we observe *Nestin*^GFP+^ cell network changes within the myenteric plexus, suggesting that the reduction in myenteric iEAN numbers is persistent in this model of infection (Figure S1H). Given the significant reduction in ileal and colonic myenteric neurons resulting from a single oral *spiB* infection, we asked whether subsequent pathogenic infection would result in exacerbated neuron loss. Following clearance of *spiB* we infected mice with wild-type *Salmonella*, after which we observed no additional loss of enteric neurons, indicating a possible restructuring or adaptation of tissue cells preventing further damage (Figure 1H). The above results demonstrate that infections with enteric bacterial and protozoan pathogens have the capacity to induce prolonged low-grade inflammation and GI dysmotility, which are associated with a rapid and long-lasting loss of iEANs.

### Enteric infections cause subtype-specific and caspase 11-dependent neuronal death

iEAN comprise a numerous and heterogeneous population of neurons that monitor and respond to various environmental cues, including mechanical stretch and luminal metabolites (Furness et al., 2013; Mayer, 2011). We first asked whether the observed loss of iEANs could be explained by a decrease of ELAVL3/4 protein expression. To address this question interbred a pan-neuronal Cre reporter mouse, *Snap25*^Cre^, with *Rpl22*^lsl-HA^ (RiboTag) mice (Sanz et al., 2009), which express a hemagglutinin (HA)–tagged ribosomal subunit 22. Immunofluorescence (IF) analysis of HA+ cells in the myenteric plexus confirmed the significant loss of iEANs as initially determined by ANNA-1 staining (Figure 2A, Figure S2A)(Muller et al., 2019). It has been reported that in cases of overt intestinal inflammation, such as colitis, there is indiscriminate loss of iEANs (Mawe, 2015). To investigate whether specific neuronal subsets were preferentially lost after enteric infection, we used confocal IF imaging to evaluate the impact on excitatory and inhibitory neuronal–cell populations (Zeisel et al., 2018). Imaging of the ileum myenteric plexus from infected and uninfected *Slc17a6*^td-Tomato^ (VGLUT2) reporter mice revealed a significant reduction in total number and percentage of excitatory VGLUT2+ neurons (Figure 2B). In contrast, quantification of inhibitory neuronal NOS+ and somatostatin (SST)+ iEAN populations revealed no changes post infection (Figure 2C, D). These data suggest a specific loss of excitatory neuronal subsets post infection, resulting in changes in the neurochemical representation of iEANs, and may provide an explanation for functional changes in GI motility observed after infection.

**Figure 2:**
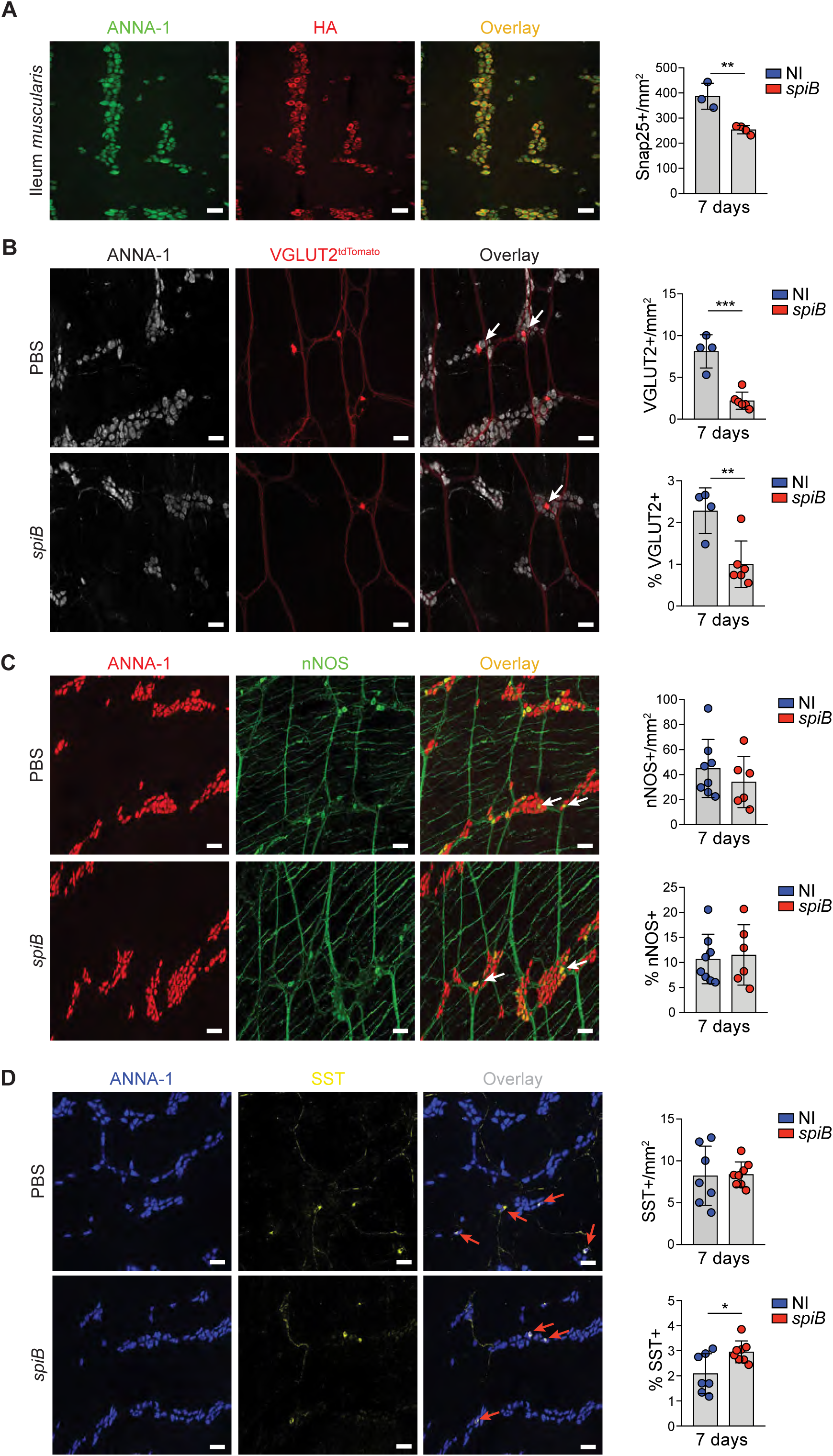
Enteric infections cause subtype-specific neuronal loss. (A) Left: Representative confocal immunofluorescence images of *Snap25*^RiboTag^ ileum myenteric plexus stained with anti-ANNA-1 (red) and anti-HA (green). Right: Quantification of the number of HA+ neurons per mm^2^ on day 7 post oral gavage with PBS (non-infected, NI) or with 10^9^ CFU of *Salmonella spiB*. (B-D) Mice were orally gavaged PBS or 10^9^ CFU of *Salmonella spiB*, and analyzed 7 days post infection. (B) Left: Representative confocal immunofluorescence image of VGLUT2^td-Tomato^ ileum myenteric plexus stained with anti-ANNA-1 (grey). Right: Quantification of the number of VGLUT2+ neurons per mm^2^ and as a percentage of ANNA-1+ neurons. (C) Left: Representative confocal immunofluorescence image of C57BL/6J ileum myenteric plexus stained with anti-ANNA-1 (red) and anti-neuronal nitric oxide synthase (nNOS) (green). Right: Quantification of the number of nNOS+ neurons per mm^2^ and as a percentage of ANNA-1+ neurons. (D) Left: Representative confocal immunofluorescence image of C57BL/6J ileum myenteric plexus stained with anti-ANNA-1 (blue) and anti-somatostatin (SST) (yellow). Right: Quantification of the number of SST+ neurons per mm^2^ and as a percentage of ANNA-1+ neurons. Data are representative of at least 3 mice per condition. At least 2 independent experiments were performed. Data were analyzed by unpaired Student’s t-test and are shown as average ± SEM; *p ≤ 0.05, **p ≤ 0.01, ***p ≤ 0.001.

To gain insights into a possible mechanism involved in infection-induced iEAN cell death, we used neuronal–specific cell–sorting independent translating ribosomal affinity purification (TRAP) (Heiman et al., 2014; Muller et al., 2019). We compared iEANs with extrinsic neurons in the nodose ganglia using *Snap25*^RiboTag^ mice, which allow neuronal-specific immunoprecipitation of actively translated mRNA. Expression of HA–tagged ribosomes was confirmed in neurons of the myenteric plexus and nodose ganglion of *Snap25*^RiboTag^ mice (Figure 2A, Figure S3A). RNA sequencing analysis of immunoprecipitated intact mRNA bound to HA–tagged ribosomes (TRAP-seq) revealed an iEAN–specific enrichment in genes encoding neuropeptides, such as neuromedin (*Nmu*), as well as components of the inflammasome pathway including *Nlrp6*, *Pycard*, *Casp1*, and *Casp11* (Figure 3A). These data indicate that iEANs possess the machinery for engaging a caspase 1/11–mediated cell death. To visualize the pattern of *Nlrp6* expression by iEANs, we performed fluorescence *in situ* hybridization (FISH) on ileum and colon sections using RNAscope® probes specific to *Elavl4* (pan-neuronal) and *Nlrp6*. We observed dense localization of *Nlrp6* transcripts in the epithelium of both the ileum and colon, similar to what has previously been reported (Levy et al., 2015). In addition, we visualized *Nlrp6* transcripts in areas of *Elavl4*– expressing cells in the *muscularis* layer, supporting the expression of NLRP6 by myenteric neurons (Figure 3B). IF staining for the inflammasome adaptor ASC (PYCARD) further confirmed the expression of this inflammasome component by iEANs of the ileum of both naïve and *spiB*– infected mice (Figure 3C, D).

**Figure 3:**
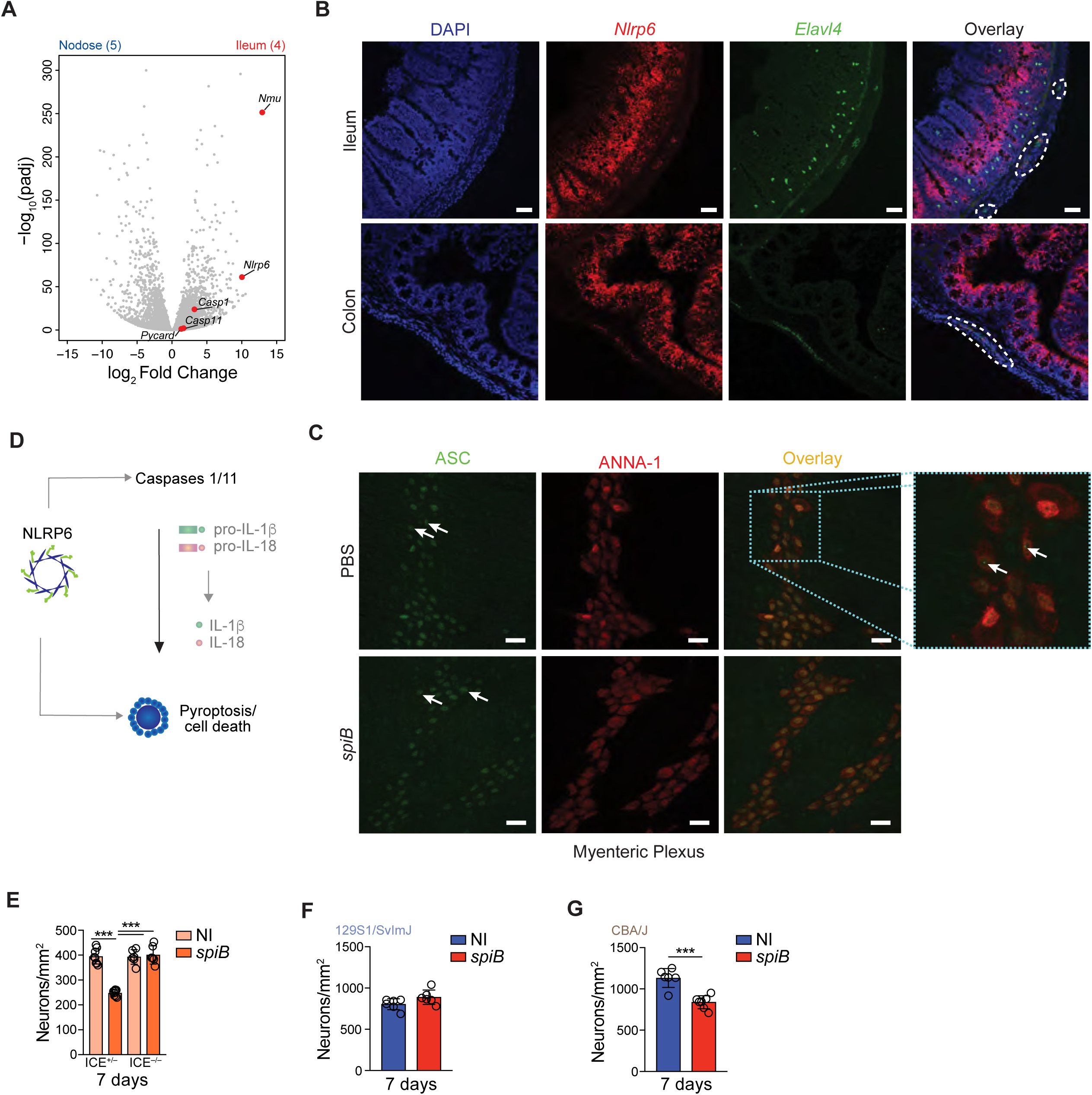
Post-infectious iEAN cell death is dependent on components of the inflammasome pathway. (A) Volcano plot of differentially expressed genes of ribosome-associated transcripts from neurons of the nodose ganglion (NG) and ileum myenteric iEANs isolated from *Snap25*^RiboTag^ mice. Grey dots highlight all genes analyzed by differential expression analysis. Red dots highlight genes that are significantly differentially expressed in ileum iEANs as compared to the NG. Sample numbers are indicated in parentheses. (B) Fluorescence *in situ* hybridization RNAScope® analysis of 15 µm sections of fresh frozen ileum and colon tissue with probes for *Nrlp6* (inflammasome component) and *Elavl4* (pan-neuronal). iEANs are outlined with a dashed line. Scale bars = 50 µm. Images representative of at least n=2 per group. (C) Immunofluorescence staining of myenteric plexus iEANs from the ileum of C57BL/6J mice using anti-ANNA-1 (red) and anti-PYCARD (ASC, green) antibodies. Samples obtained at 6 hours post oral gavage of PBS or 10^9^ CFU of *Salmonella spiB*. Arrows indicate ASC “specks”; inset shows ImageJ zoom of the indicated panel in PBS-treated mice. Scale bars = 25 µm. Images representative of at least n=5 per group. (D) Simplified schematic of the NLRP6 inflammasome pathway. (E) Neuronal quantification as assessed by immunofluorescence staining (ANNA-1) in the myenteric plexus of ileal segments on day 7 post *spiB* infection of *Casp1*^-/-^ *Casp11*^-/-^ (Interferon-converting enzyme, ICE) double-knockout (ICE^-/-^) mice or heterozygous controls (ICE^+/-^). (F, G) Neuronal quantification as assessed by immunofluorescence staining (ANNA-1) in the myenteric plexus of colonic segments on day 7 post *spiB* infection of 129S1/SvImJ mice and (F) CBA/J mice. Unless otherwise indicated, data are representative of 3-6 mice per condition. At least 2 independent experiments were performed. Data were analyzed by unpaired Student’s t-test or ANOVA with Bonferroni correction and are shown as average ± SEM; n.s. - not significant, *p ≤ 0.05, **p ≤ 0.01, ***p ≤ 0.001, ****p≤ 0.0001.

To address the role of caspase–mediated cell death in iEAN loss during infection, we systemically administered a pan caspase inhibitor (zVAD-FMK) to mice infected with *spiB*, which reduced infection-induced iEAN loss (Figure S3B). We directly addressed the role of caspases 1 and 11 in infection–induced iEAN loss by infecting caspase 1/11 (ICE)–deficient or haplosufficient (ICE^+/^) mice (Kuida et al., 1995) with *spiB*. While ICE^+/–^ mice exhibited pronounced iEAN loss in the ileal MP 7 days post–infection, *spiB*-infected ICE^−/−^ mice were completely protected from neuronal loss, despite similar bacterial clearance patterns (Figure 3D, E, see CFU summary in Figure S6A). We further dissected the specific role caspases 1 and 11 in neuronal cell death by utilizing the 129 mouse strain, which carries an inactivating mutation in the *Casp11* locus (human *Casp4*) (Kenneth et al., 2012; Vanden Berghe et al., 2015) (Figure S3C-F). 129 mice infected with *spiB* exhibited no loss of colonic iEAN numbers as compared to non-infected controls. (Figure 3F). This iEAN protection was independent of the ability of 129 mice to survive WT *Salmonella* infection due to expression of functional *Nramp* (Brown et al., 2013), as we found that CBA/J mice, which also express a functional *Nramp*, but do not carry a *Casp11* mutation, exhibited significant neuronal loss when infected with *spiB*, despite similar clearance of the bacteria (Figure 3G, Figure S3C-F, and *data not shown*). These data implicate caspase 11–mediated cell death as a main mechanism involved in iEAN loss following *Salmonella* infection.

### MMs respond to luminal infection to limit neuronal damage

Tissue–resident MMs, the most abundant immune cell population in the myenteric plexus, are closely juxtaposed to iEANs, and their presence has been linked to normal functioning of these neurons during homeostasis (De Schepper et al., 2018; Gabanyi et al., 2016; Muller et al., 2014). Our previous observations using both cell-sorting– and TRAP–based approaches, suggested that MM preferentially express tissue-protective and wound healing genes, such as *Retnla* (encoding Fizz1), *Mrc1*, *Cd163* and *Il10* at steady state. This gene signature was reinforced early (2 h) after oral exposure to *spiB*, with an upregulation of additional tissue-protective genes such as *Arg1* and *Chi3I3* (encoding Ym1) (Gabanyi et al., 2016). To investigate a possible role of MMs in infection– induced iEAN damage, we first depleted MMs using an antibody blocking colony stimulating factor 1 receptor (CSF1R)–signaling, AFS98 (α-CSF1R). We used a dose that preferentially depletes MMs over lamina propria macrophages, due to the fact that MMs express higher levels of CSF1R (Muller et al., 2014) (Figure 4A, Figure S4A, B, and Figure S6B). Continuous α-CSF1R-mediated MM depletion did not impact iEAN numbers in naïve mice. However, MM depletion resulted in an enhanced iEAN loss in mice infected with *spiB* when compared to mice treated with isotype control antibody, despite similar bacterial load in both conditions (Figure 4B, Figure S6B). These results indicate that while short–term depletion of MMs does not impact iEAN survival in the unperturbed state, MMs may play an iEAN–protective role in the context of enteric infection.

**Figure 4:**
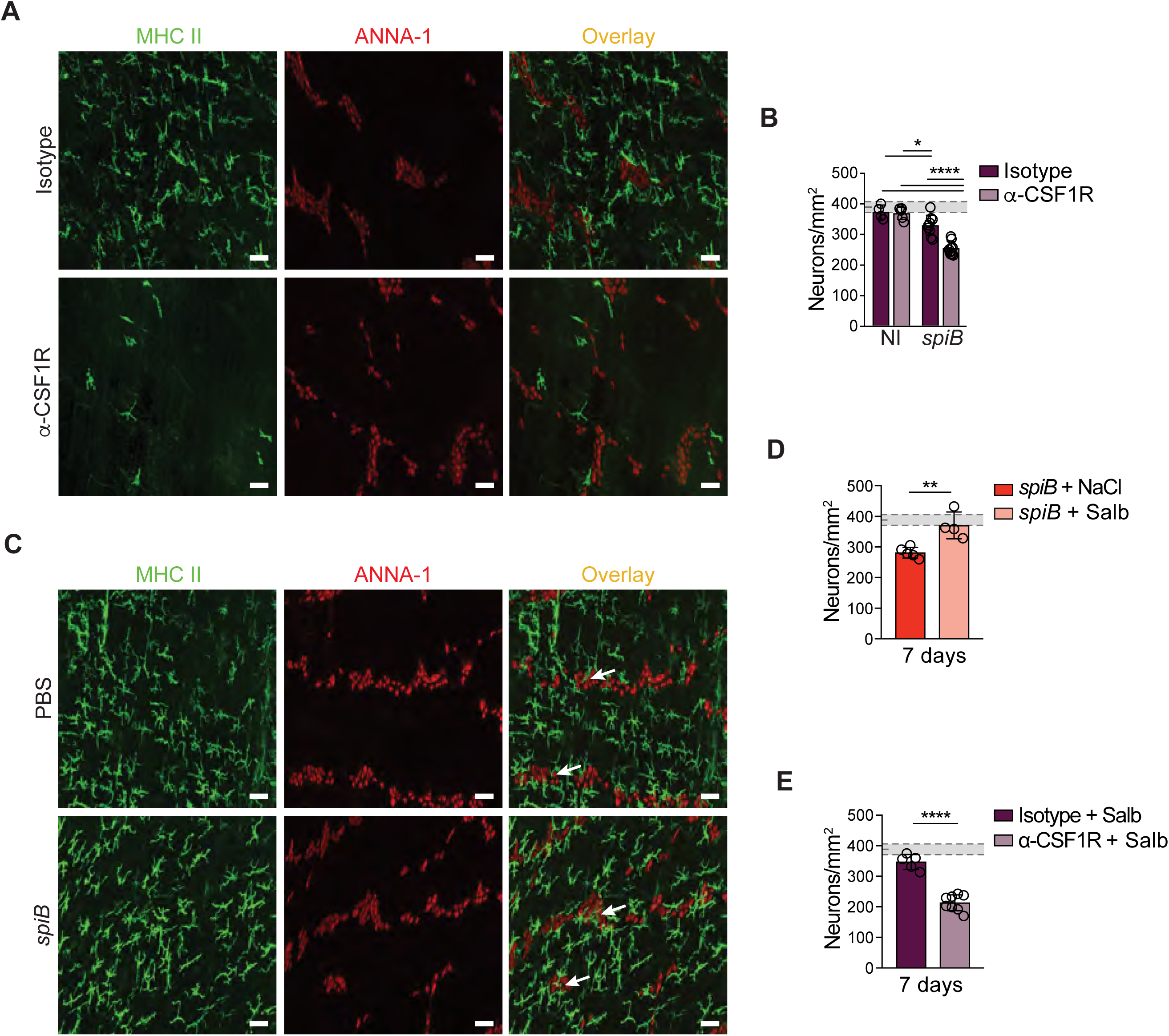
Tissue-resident MMs respond to luminal infection to limit neuronal loss. (A, B) C57BL/6J mice orally gavaged PBS or 10^9^ CFU of *Salmonella spiB* were treated with anti-CSF1R or IgG isotype control antibodies by intraperitoneal injections twice over the course of a 7-day period. (A) Representative confocal immunofluorescence staining of MM and iEAN of the ileum myenteric plexus region stained with anti-MHC-II (green) and anti-ANNA-1 (red). Images representative of at least n=5 per condition; (B) Neuronal quantification as assessed by immunofluorescence staining (ANNA-1) of ileum myenteric plexus of injection-habituated non-infected and *spiB*-infected C57BL/6J mice receiving anti-CSF1R or IgG isotype control antibodies; (C) Representative confocal immunofluorescence images from the ileum myenteric plexus of C57BL/6J mice using anti-MHC II (green) and anti-ANNA-1 (red). Samples obtained at 3 hours post oral gavage of PBS or *spiB*. (D) Neuronal quantification as assessed by immunofluorescence staining (ANNA-1) in the myenteric plexus of C57BL/6J mice continuously receiving either vehicle (NaCl) or salbutamol sulfate delivered by subcutaneous osmotic pumps for 14 days, and orally gavaged PBS or 10^9^ CFU of *Salmonella spiB* on day 7 post-surgical pump implantation. Samples obtained on day 7 post infection. (E) Neuronal quantification as assessed by immunofluorescence staining (ANNA-1) in the myenteric plexus of C57BL/6J mice receiving salbutamol delivered by subcutaneous osmotic pumps for 14 days. Mice were gavaged PBS or *spiB* on day 7 post-surgical pump implantation and received anti-CSF1R or IgG isotype control antibodies over the course infection. Samples obtained at day 7 post infection. (B, D-E) Shaded area bounded by dashed lines indicates mean day 7 iEAN numbers +/- SEM of all control C57BL6/J mice analyzed in Figure 1F. Data are representative of at least n=3 per condition. At least 2 independent experiments were performed. Data were analyzed by unpaired Student’s t-test and are shown as average ± SEM; *p ≤ 0.05, **p ≤ 0.01, ***p ≤ 0.001, ****p≤ 0.0001.

Next, we used confocal imaging to investigate MM dynamics early upon infection, which revealed a continuing presence and intercalation into ganglia of the myenteric plexus (Figure 4C, Figure S4C). To assess whether inflammatory monocytes recruited from the circulation are involved in infection-induced neuronal damage, we used CCR2^−/−^ mice, which exhibit an impairment in mobilizing inflammatory monocytes into tissues (Boring et al., 1997). CCR2^−/−^ mice failed to clear *spiB* infection, likely reflecting a reduced resistance in the mucosal layer (Dunay et al., 2008), and showed mild acceleration, rather than a delay in GITT at 17 dpi (Figure S4D, E). Nonetheless, CCR2^−/−^ mice showed similar iEAN loss as WT control mice (~25%), suggesting that the persistent luminal pathogen load does not cause an increase in neuronal damage (Figure S4F). These data also highlight that intestinal macrophages recently differentiated from circulating monocyte precursors may contribute to *spiB* clearance mechanisms, but are not required for post–infectious iEAN cell death. The above results suggest a rapid response of resident MMs to luminal pathogenic stimulation; while MMs are likely not essential for resistance mechanisms, they appear to prevent excessive neuronal damage.

### β2-AR signaling in MMs constrains infection–induced inflammation and neuronal loss

We previously reported that the tissue-protective gene-expression profile of MMs was further imprinted following enteric infection via β_2_ adrenergic receptors (β_2_-ARs)(Gabanyi et al., 2016). We investigated the capacity of β_2_-AR signaling to mediate tissue protection by coupling pharmacological and genetic modulation of this pathway. First, C57BL/6 mice were infected with *spiB* while being treated with a selective β_2_-AR agonist (salbutamol) which was delivered continuously via subcutaneously implanted osmotic pumps for 14 days. We did not observe differences in the pathogen load in mice treated with salbutamol (Figure S6C). However, administration of a salbutamol protected mice from neuronal loss post infection (Figure 4D). To directly assess the role of MMs in the observed salbutamol–mediated protection from neuronal death post infection, we then depleted MMs using α-CSF1R in mice receiving continuous salbutamol treatment. While we observed a rescue of iEAN death in mice treated with IgG isotype control antibody, MM depletion led to a loss of salbutamol-mediated iEAN protection (Figure 4E). Taken together, these data suggest that MMs are critical for β_2_-AR–mediated iEAN protection.

We next addressed whether this adrenergic signaling in macrophages plays a role in infection– induced enteric neuronal damage utilizing a genetic approach. We interbred mice carrying Cre recombinase under the myeloid *Lyz2* promoter (LysM^Cre^) with *Adrb2^flox/flox^* mice (Hinoi et al., 2008). The specificity of Cre-targeting of macrophages in the intestinal *muscularis* was confirmed by immunohistochemistry using HA as a fate mapping reporter (Figure 5A). We did not detect infiltrating neutrophils, also targeted by *Lyz2*, in the intestinal *muscularis* at steady state or early post infection, suggesting that neutrophil-β_2_-AR signaling does not play a major role in this model(*data not shown*). Additionally, we did not observe differences in iEAN numbers or GITT in LysM^Δ^*^Adrb2^* as compared to Cre^−^ littermate control mice during homeostasis (Figure S5A, B). Upon *spiB* infection, LysM^Δ^*^Adrb2^* also exhibited a similar pathogen clearance (Figure 5B). However, we found that LysM^Δ^*^Adrb2^* mice had further increased fecal Lcn-2 levels following *spiB* infection as compared to Cre^−^ littermates, suggesting a role for myeloid-specific β_2_-AR signaling in the regulation of infection–induced inflammation (Figure 5C). This enhanced intestinal inflammation was accompanied by further prolonged GITT, which persisted long-term post *spiB* clearance, an increased loss of myenteric neurons and enhanced alterations in ileal ring contractility (Figure 5D, E, Figure S5C). To complement these results, and to address whether MMs themselves require intact β_2_-AR signaling to limit neuronal loss, we infected LysM^Δ^*^Adrb2^* mice and Cre^−^ littermate controls receiving continuous salbutamol treatment. Similar to what we observed in wild-type C57BL/6 mice, salbutamol treatment in Cre^−^ littermate controls carrying an intact β_2_-AR signaling in the myeloid compartment resulted in prevention of infection-induced neuronal loss. Conversely, salbutamol–treated LysM^Δ^*^Adrb2^* mice still displayed significant iEAN loss (Figure 5F). These data establish a role for β2AR signaling in intestinal macrophages that contributes to a neuroprotective program post enteric infection.

**Figure 5:**
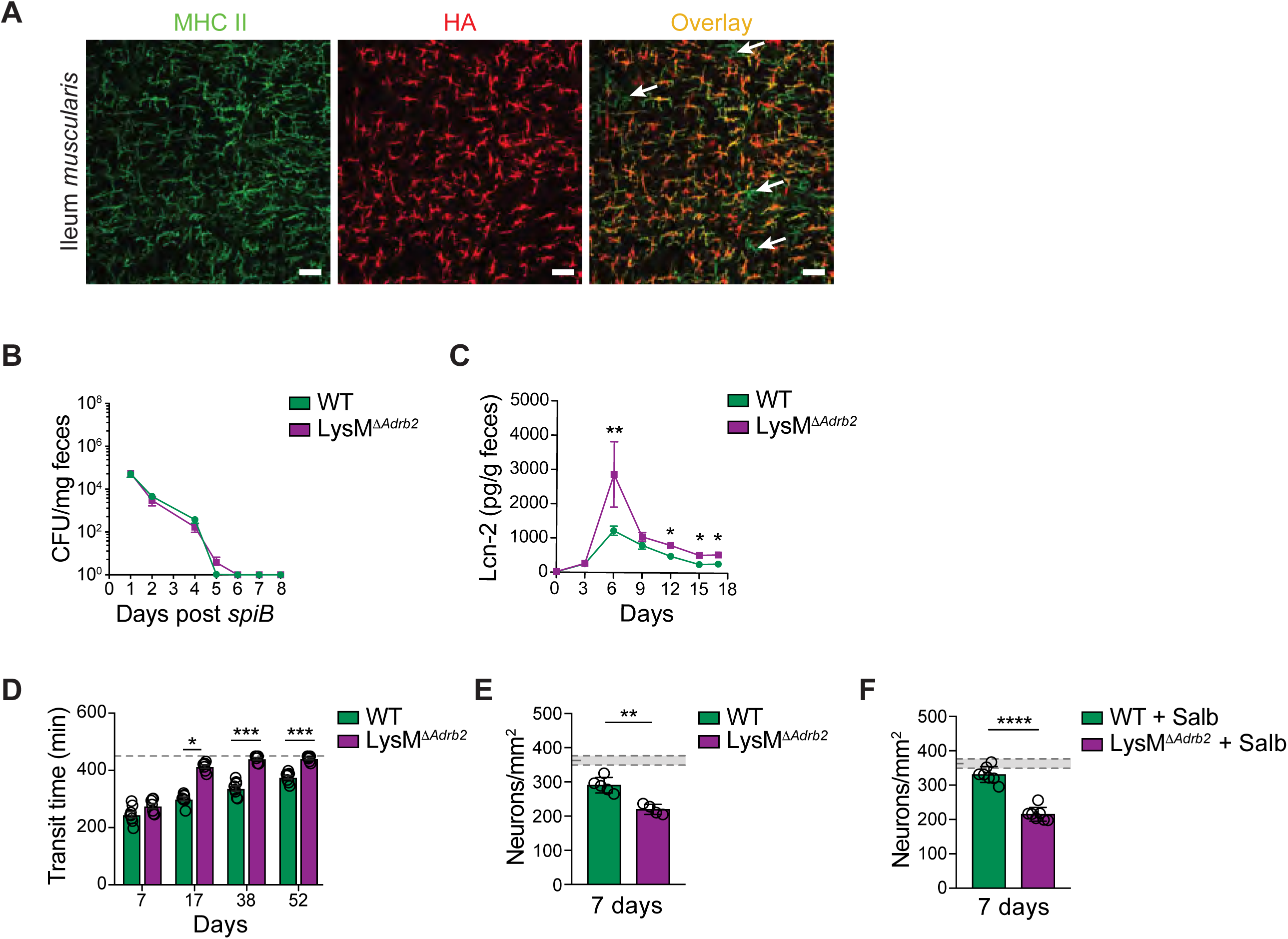
Loss of β_2_–AR signaling in MMs exacerbates neuronal damage during enteric infection. (A) Immunofluorescence staining of MMs from the ileum of LysM^RiboTag^ mice using anti-MHC II (green) and anti-hemagglutinin (HA, red) antibodies. White arrows indicate MHC II+ HA-macrophages. Scale bars = 50 µm. Images representative of n=2 mice per group. (B-F) LysM^Δ*Adrb2*^ and wild-type (WT) littermate control mice were orally infected with 10^9^ CFU of *Salmonella spiB*. (B) Quantification of fecal colony forming units (CFU) on the indicated days post infection (C) Quantification of fecal lipocalin-2 (Lcn-2) on the indicated days post infection. (D) Gastrointestinal motility activity as assessed by total gastrointestinal transit time; experiments were ended at 450 min (dashed line); (E, F) Neuronal quantification as assessed by immunofluorescence staining (ANNA-1) in the myenteric plexus of ileal segments on day 7 post infection with *spiB* of (E) LysM*^ΔAdrb2^* and wild-type (WT) littermate control mice; shaded area bounded by dashed lines indicates mean day 7 iEAN numbers +/- SEM of non-infected LysM^Δ*Adrb2*^ and WT littermate control mice (Figure S5B); (F) LysM*^ΔAdrb2^* and wild-type (WT) littermate control mice receiving 400 µg/day salbutamol sulfate delivered by subcutaneous osmotic pumps for 14 days. Mice were infected with *spiB* on day 7 post-surgical pump implantation; shaded area bounded by dashed lines indicates mean day 7 iEAN numbers +/- SEM of non-infected LysM^Δ*Adrb2*^ and WT littermate control mice (Figure S5B). Unless otherwise indicated, data are representative of 3-6 mice per condition. At least 2 independent experiments were performed. Data were analyzed by unpaired Student’s t-test or ANOVA with Bonferroni correction and are shown as average ± SEM; *p ≤ 0.05, **p ≤ 0.01, ***p ≤ 0.001, ****p≤ 0.0001.

It did not escape our attention that mice i.p.-injected multiple times with IgG isotype control antibodies (as in MM depletion experiments) or mice anesthetized with isoflurane (as in control pump experiments) did not lose usual (~25%) iEAN numbers post *spiB* infection, suggesting that stress–induced local or systemic catecholamine release could trigger the same protective pathway in MMs. Indeed, a short pulse of isoflurane anesthesia or i.p. injections of PBS or IgG isotype antibody was sufficient to prevent most iEAN cell death following *spiB* infection, suggesting that stress signals that potentiate β_2_AR signaling may help in preventing post–infectious iEAN loss (Figure S5D, E). These results further support a role for catecholamines and β_2_AR signaling in MMs in the regulation of inflammation–induced iEAN loss.

### An adrenergic-arginase-polyamine axis in MMs limits infection–induced neuronal loss

To dissect possible mechanisms by which MM β_2_-AR signaling is involved in preventing excessive tissue damage post–infection, we analyzed LysM^Δ^*^Adrb2^* MM gene expression profiles. MMs sorted from Cre^−^ littermate controls responded to *spiB* infection by upregulating tissue protective genes upon infection, but we did not observe this upregulation in MMs sorted from infected LysM^Δ^*^Adrb2^* mice (Figure 6A). Arginase 1 is known to mediate the production of neuroprotective polyamines such as spermine (Cai et al., 2002), which in turn were described to suppress NRLP6– caspase1/11 inflammasome activation (Levy et al., 2015). To investigate the participation of polyamines in iEAN cell death, we supplemented the drinking water of mice infected with *spiB* with Spermine or with Difluoromethylornithine (DFMO), which inhibits polyamine biosynthesis by selective and irreversible inhibition of ornithine decarboxylase 1 (ODC1)(Koomoa et al., 2013). Bacterial load and clearance patterns were similar in either treatment condition; however, mice that received DFMO exhibited enhanced neuronal loss, while those exposed to spermine showed a significant rescue of neuronal loss post *spiB* infection (Figure 6B, C, Figure S6D, E). Finally, we genetically addressed whether Arginase 1 activity in MMs was required for their protective role in infection–induced neuronal damage in the intestine by interbreeding LysM^Cre^ and *Arg1^flox/flox^* mice (El Kasmi et al., 2008). Upon *spiB* infection, LysM*^ΔArg1^* also showed similar pathogen load and clearance patterns (Figure S6F). However, loss of Arginase 1 in the myeloid compartment significantly heightened fecal Lcn-2 levels and iEAN loss following *spiB* infection (Figure 6D). Together, these results point to a functional role for β_2_AR–Arginase 1-polyamines axis in MM–mediated tissue protective program in limiting infection–induced enteric neuronal cell death.

**Figure 6:**
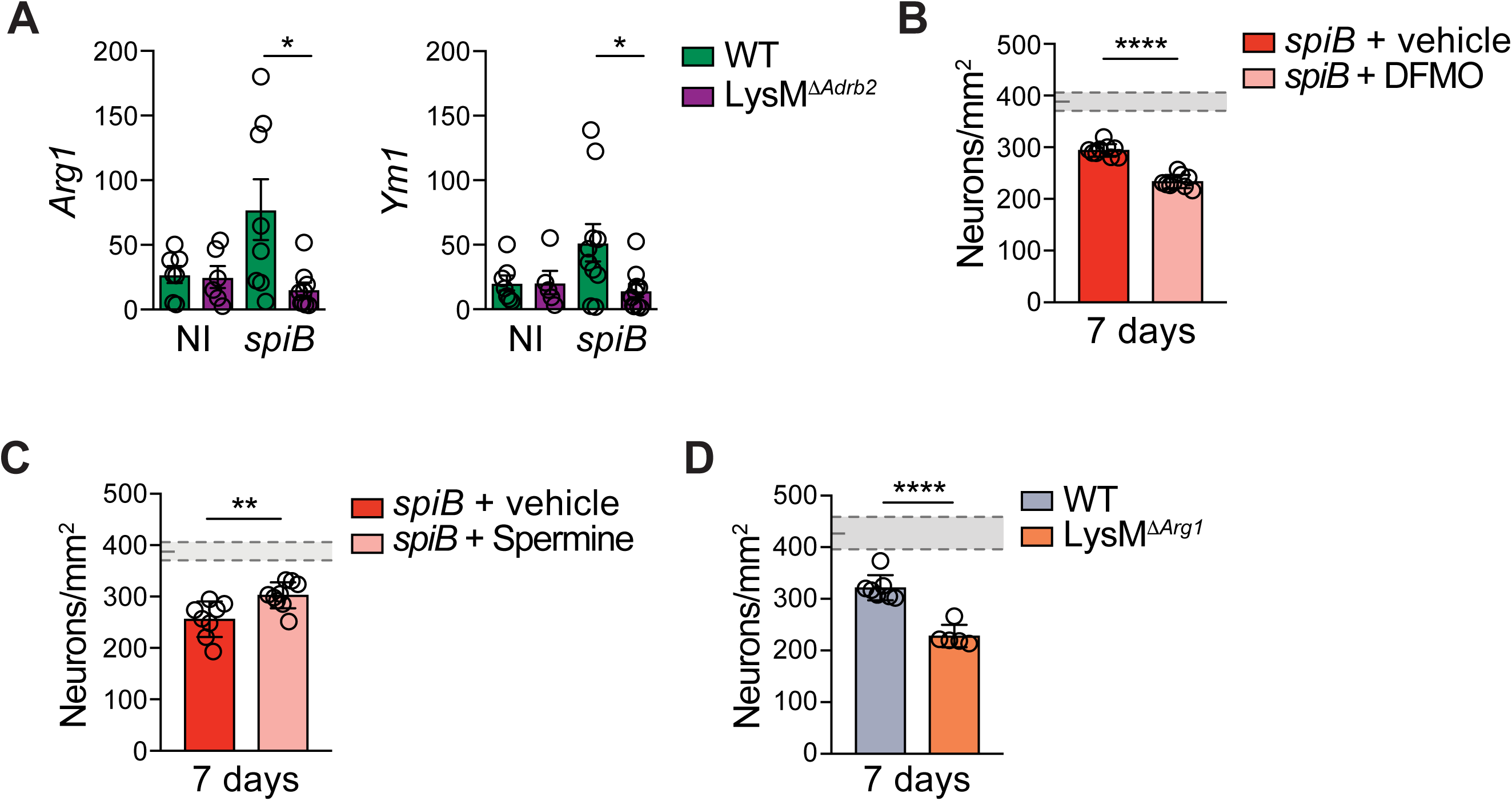
An adrenergic-arginase-polyamine axis in MMs limits infection-induced neuronal loss. (A) mRNA expression analysis of the indicated genes by quantitative real-time PCR in LysM^Δ*Adrb2*^ and wild-type (WT) littermate control mice non-infected (NI) or 2 hours post infection with 10^9^ CFU of *Salmonella spiB*. (B-D) Neuronal quantification as assessed by immunofluorescence staining (ANNA-1) in the myenteric plexus of ileal segments on day 7 post infection with *spiB* of (B) C57BL/6J mice with access to regular drinking water or water supplemented with difluoromethylornithine (DFMO) over the course of infection starting 5 days prior to infection; shaded area bounded by dashed lines indicates mean day 7 iEAN numbers +/- SEM of all control C57BL6/J mice (C) C57BL/6J mice receiving regular drinking water or water supplemented with Spermine over the course of infection starting 3 days prior to infection; shaded area bounded by dashed lines indicates mean day 7 iEAN numbers +/- SEM of all control C57BL6/J mice (D) LysM^Δ*Arg1*^ mice and wild-type (WT) littermate controls; shaded area bounded by dashed lines indicates mean day 7 iEAN numbers +/- SEM of non-infected LysM^Δ*Arg1*^ and WT littermate control mice (Figure S5C). Data are representative of 3-6 mice per condition. At least 2 independent experiments were performed. Data were analyzed by unpaired Student’s t-test or ANOVA with Bonferroni correction and are shown as average ± SEM; *p ≤ 0.05, **p ≤ 0.01, ***p ≤ 0.001, ****p≤ 0.0001.

## Discussion

### Inflammation–induced neuronal damage

Overt inflammation in response to infection has the capacity to result in chronic immunopathology. In the GI tract, bacterial, viral and protozoan infections are associated with intestinal neuropathy and the development of post-infectious IBS (Beatty et al., 2014; Holschneider et al., 2011; Ohman and Simren, 2010; White et al., 2018). However, the understanding of underlying mechanisms involved in this process is largely lacking. Here, we provide evidence that *muscularis* macrophages located in close proximity to enteric neurons prevent infection– or inflammation– induced tissue damage; perhaps an analogous function to that described for microglia in the CNS (Davalos et al., 2005; El Khoury et al., 2007; Nimmerjahn et al., 2005; Wang et al., 2015).

Most mechanisms proposed for inflammation–induced neuronal damage thus far are related to CNS neuroinflammation, including Alzheimer’s disease (AD), Parkinson’s disease (PD), amyotrophic lateral sclerosis (ALS), multiple sclerosis (MS), which have been associated with impaired neurological function (Aguzzi et al., 2013; Colonna and Butovsky, 2017). Although certain inflammatory processes are essential for neuronal function and tissue physiology, including neuronal pruning and tissue repair (Salter and Stevens, 2017), excessive inflammation often result in the production of neurotoxic factors such as interleukin 1 beta (IL-1β), tumor necrosis factor alpha (TNF-α), nitric oxide (NO) and reactive oxygen species (ROS) (Crotti and Glass, 2015). Additionally, neurotropic viruses such as West Nile virus, may directly induce neuronal death and encephalitis (Samuel et al., 2007; Shrestha et al., 2003); indirectly, neurotropic virus infections, including Zika virus, can activate CD8^+^ T– or NK cell–mediated killing of infected neurons, including EANs (White et al., 2018). While some of these factors could play a role in the models examined here, our studies rather point to NLRP6 ligands, or alternative triggers of caspase 11 cleavage, as mediators of infection–induced iEAN cell death. These results corroborate previous work describing a protective phenotype in *Pycard*^−/−^ mice, which did not lose enteric neurons following DNBS-induced colitis, a process that was also found to be NLPR3-independent (Gulbransen et al., 2012). Unlike the indiscriminate iEAN loss previously observed in chemically–induced murine colitis models (Linden et al., 2005), our studies indicate infection-induced location- and subset-specific loss of excitatory iEANs, and a preservation of the main inhibitory populations. These data warrant further study and additional subset-specific analysis of EANs in the steady state or under pathological conditions, although genetic tools for these approaches are still lacking. Nevertheless, we speculate that the mechanism of neuronal cell death involving components of the inflammasome machinery put forward here in the context of enteric infection may be engaged in additional contexts. For instance, it is possible that iEAN– mediated production of IL-1β or IL-18, via caspase 1/11 activation, participate in pathogen resistance mechanisms (R. Flavell, personal communication). Furthermore, it remains to be determined whether neurodegenerative processes in the CNS (Song and Colonna, 2018) also involve this or an analogous inflammasome-caspase axis.

### Monocyte-macrophage lineage in neuronal damage

Studies utilizing murine models and human tissue have uncovered a potential role of monocytes, brain endothelial cells, astrocytes, macrophages and microglia in the induction, or regulation of inflammation–induced neuropathy, CNS damage, and subsequent changes in animal behavior (Aguzzi et al., 2013; Colonna and Butovsky, 2017; D’Mello et al., 2013; Habbas et al., 2015). Tissue resident macrophage populations in the brain, and in particular microglia within the parenchyma, express PRRs and are believed to contribute significantly to dendritic spine remodeling (Parkhurst et al., 2013; Song and Colonna, 2018). The balance between the production of pro-inflammatory mediators and tissue protective factors by macrophage populations is likely to determine the pathological versus reparative role of these cells. In a model mimicking CNS virus infection, depletion of CX3CR1+ cells (including monocyte-macrophage lineage), but not of CNS resident macrophages (including microglia (Parkhurst et al., 2013)), abrogated poly(I:C)-mediated dendritic spine loss, thereby implicating monocytes or monocyte-derived cells, rather than CNS-resident macrophages, in a TLR signaling and TNF-*α* production– mediated neuronal damage (Garré et al., 2017). The synergistic influence of TNF-*α* on type Iinterferons could play an important role, as the latter are also linked to neuronal damage, in disrupted synaptic connectivity and cognitive impairment during viral infections (Venkatesh et al., 2013). In the intestine under stress or inflammatory conditions, activation of enteric glia (gliosis) have also been linked to neuronal damage (Brown et al., 2016; Gulbransen et al., 2012; Mayo et al., 2014), although very little is known on the relevance of glial-immune cell interactions in the intestinal tract. It is therefore plausible that surrounding enteric glia, or MMs, induce neuronal damage as previously attributed to microglia in the brain (Jin et al., 2016; Lehnardt, 2010).

In contrast to pathological roles described above, both glial cells and tissue macrophages are also appreciated to prevent inflammation–induced damaged in several different contexts, as well as to initiate post injury repair processes. In models of spinal cord injury, for example, incoming monocytes are thought to mediate a functional recovery via secretion of anti-inflammatory mediators (Shechter et al., 2009), although an enhancement of macrophage inflammatory markers was also shown to help CNS axonal regeneration (de Lima et al., 2012; Gensel et al., 2009). Our data favors a neuronal protective role for *muscularis*–resident macrophages via upregulation of arginase 1 and production of polyamines; an analogous observation to previous studies suggesting that “alternatively activated” macrophages promote axonal growth or regeneration after CNS injury (Cai et al., 2002; Kigerl et al., 2009; Kuo et al., 2011). While multiple studies of the CNS find a direct role for polyamines in neuro-regeneration (Mahar and Cavalli, 2018), our data suggest a potential role for these molecules in preventing neuronal cell death via inhibition of the inflammasome. This latter possibility is supported by previous studies demonstrating that microbiota–derived spermine regulates NLRP6 inflammasome signaling and intestinal epithelial cell secretion of IL-18 (Levy et al., 2015), all of which could also play tissue protective roles in the enteric infection models described here. Nonetheless, macrophages differentiated towards an “alternatively activated” phenotype could also negatively contribute to gut homeostasis during the resolution of inflammatory processes, for instance by enhancing fibrosis (Xue et al., 2015). Finally, because we previously demonstrated that macrophages located in the mucosal layer do not show an immediate upregulation of tissue protective genes upon enteric infection, an anatomically segregated model in which lamina propria macrophages help pathogen resistance and distally–located MMs boost tolerance is conceivable.

### Sensing and regulation of the catecholamine-β_2_-AR pathway

The nervous and immune systems are the body’s main sensory interfaces that perceive, integrate and respond to environmental challenges. Both systems depend on cell-to-cell contacts and on soluble molecules that act on proximal or distant target cells, which include neurons and immune cells themselves (Besedovsky et al., 1983). In addition to the anti-inflammatory reflex, the involvement of catecholaminergic neurons in anti-inflammatory responses has been demonstrated in different tissues, and are attributed to the broad sympathetic innervation in peripheral lymph nodes associated with expression of β_2_-AR in immune cells (Pavlov et al., 2018). Our findings also suggest an intricated interplay between iEANs, macrophages and eEANs, particularly the extrinsic sympathetic ganglia, adding functional significance to observations of interactions between barrier tissue resident macrophages and peripheral neurons (Balmer et al., 2014; Gabanyi et al., 2016; Muller et al., 2014). The source of catecholamines may be context dependent: enteric infections, including *Salmonella*, may result in the activation of extrinsic sympathetic neurons located in the celiac-superior mesenteric ganglion (Gabanyi et al., 2016); under conditions of systemic stress these signals could originate from the adrenal glands, a possibility also supported by our findings. Both of these pathways nevertheless require neural input, although the circuitry connecting luminal sensing to eEAN activation (and a possible involvement of the CNS), and how luminal stimulations can regulate it, remain to be determined (Muller et al., 2019; Pruss et al., 2017). Current literature suggests that the epithelium layer works as a primary site of sensory integration. In particular, enteroendocrine cells are uniquely tuned to sense and respond to luminal changes, since they are directly connected with neurons innervating the small intestine and colon, forming functional synaptic connections with sensory neurons (Bohorquez et al., 2015; Kaelberer et al., 2018). A non-mutually exclusive possibility is that neuronal processes reaching the intestinal *villi* directly sense microbial or stress signals (Chiu et al., 2013), either directly modulating iEAN survival or initialing changes in eEANs. However, the components of luminal sensing still require further studies specifically targeting stress– or pathogen– recognition receptors or other potential direct sensing channels or receptors in EANs (Veiga-Fernandes and Mucida, 2016). Regardless of the initial components, our data identify a novel mechanism of neuronal cell–death and point to a role for tissue–resident MMs in the regulation of neuronal damage. Such intra-tissue adaptation of immune cells appears to help the maintenance of a proper balance between resistance and tolerance and these findings could have ramifications in additional tissues or contexts.

## Author Contributions

P.A.M initiated, designed, performed, analyzed and helped supervising the research and wrote the manuscript; F.M. designed, performed and analyzed the research, and wrote the manuscript; C.G. designed and analyzed experiments, performed enteric infection, macrophage dynamics and EAN-loss experiments, and helped writing the manuscript; I.G. initiated and made initial observations of EAN loss post enteric infections; Z.K. performed experiments; D.B. helped with EAN quantification and protozoan infections; D.M. conceived, designed and supervised the research, and wrote the manuscript.

## Acknowledgments

We thank all Mucida Lab members, past and present, for assistance in experiments, fruitful discussions and critical reading of the manuscript. We also thank Michel Nussenzweig, Gabriel Victora (Rockefeller University, RU) and Juan Lafaille (NYU) and their respective lab members for fruitful discussions and suggestions. Aneta Rogoz for the maintenance and genotyping of gnotobiotic mice, Telmo Catarino (Champalimaud Center, Portugal) for help with EAN quantification, Sara Gonzalez for maintenance of SPF mice, RU Bio-imaging Research Center for assistance with confocal microscopy and image analysis, RU Genomics Center for RNA sequencing and the RU employees for continuous assistance. We thank members of our laboratory for discussions and critical reading and editing of the manuscript. We thank Jeffrey Friedman (RU) for the generous use of lab equipment, S. Ackerman and P. Cohen (RU) for insightful discussion and help with DFMO experiments. We thank P. Woster (MUSC) for kindly providing DFMO, and Vanda A. Lennon (Mayo Clinic) for generously sharing anti-ANNA-1. We thank R. Gazzinelli (U. Mass.) for kindly providing *T. cruzi* and *T. gondii* strains. We also thank Yasmine Belkaid (NIH) for kindly providing *Yersinia pseudotuberculosis*, Miriam Merad (MSSM) for kindly sharing the ASF98 hybridoma cell line, Paul Frenette (Einstein) and Grigori Enikolopov (Stony Brook) for generously sharing Nestin^GFP^ mice, and Gerard Karsenty (CUMC) for kindly sharing Adrb2^flox/flox^ mice. We also thank This work was supported by NIH F31 Kirchstein Fellowship (P.A.M.), NCATS NIH UL1TR001866 (P.A.M., D.M.), Anderson Graduate Fellowship (F.M.), The Rockefeller University Women in Science Fellowship (F.M.), Philip M. Levine Fellowship (P.A.M.), Kavli Graduate Fellowship (P.A.M.), the Leona M. and Harry B. Helmsley Charitable Trust (F.M., P.A.M., D.M.), the Crohn’s & Colitis Foundation of America (D.M.), the Kenneth Rainin Foundation (D.M.), and National Institutes of Health NIH R01 DK093674 and R01 DK113375 grants (D.M.).

## Supplementary Figure legends

**Figure S1 (related to Figure 1).**

(A) Neuronal quantification as assessed by immunofluorescence staining (ANNA-1) in the myenteric plexus of ileal segments on the indicated days post exposure of C57BL/6J mice to 10^9^ CFU of heat-killed *Salmonella spiB*. (C) Neuronal quantification as assessed by immunofluorescence staining (ANNA-1) in the myenteric plexus of ileal segments C57BL/6J mice orally infected with 10^8^ CFU of *Y. pseudotuberculosis*. (D-F) C57BL/6J mice were orally infected with 5 cysts of *Toxoplasma gondii*. (D) Quantification of fecal Lcn-2 levels as assessed by ELISA on the depicted days; (E) Gastrointestinal motility as assessed by total gastrointestinal transit time on the indicated days post infection; experiments were ended at 450 min (dashed line). (F) neuronal quantification as assessed by cuprolinic blue staining in the myenteric plexus of ileal segments on the indicated days post infection. (G) Neuronal quantification as assessed by cuprolinic blue staining in the myenteric plexus of ileal segments on the indicated days post infection of C57BL/6J mice with 10^4^ parasites of *T. cruzi*. (H) Representative confocal immunofluorescence images of ileum muscularis obtained from *Nestin*^GFP^ reporter mice 7 days post oral exposure to PBS or *spiB*. Scale bar = 50 µm. Images representative of at least n=5 animals per group. Data are representative of at least 3 mice per condition. At least 2 independent experiments were performed. Data were analyzed by unpaired Student’s t-test or ANOVA with Bonferroni correction and are shown as average ± SEM; *p ≤ 0.05, **p ≤ 0.01, ***p ≤ 0.001.

**Figure S2 (related to Figure 2).**

Quantification of the number of *Snap25*^RiboTag^+ neurons as a percentage of ANNA-1+ neurons.

**Figure S3 (related to Figure 3).**

(A) Representative confocal immunofluorescence images of *Snap25*^RiboTag^ nodose ganglia stained with anti-ANNA-1 (red) and anti-HA (green). Scale bars = 50 µm. Images representative of n= 5 mice. (B) Neuronal quantification as assessed by immunofluorescence staining (ANNA-1) in the myenteric plexus of ileal segments on day 7 post *spiB* infection of C57BL/6J mice receiving vehicle (dimethyl sulfoxide, DMSO) or zVAD-FMK (150 µg/day) by daily intraperitoneal injections over the course of infection. Shaded area bounded by dashed lines indicates mean day 7 iEAN numbers +/- SEM of all control C57BL6/J mice analyzed in Figure 1F. (C-F) Analysis of *Casp11* mutations in mouse strains used. (D) 3% agarose gel for *Casp11* 5bp deletion PCR product. Mouse strains used in this work are indicated below each lane. Red dashed line indicates the size at which a potential 5bp deletion PCR product will run. (E) Sanger sequencing of *Casp11* 5bp deletion PCR product in C57BL/6J, CBA/J, and 129S1/SvImJ mice. (F) Sequence surrounding *Casp11* 5bp deletion region in C57BL/6J, CBA/J, and 129S1/SvImJ mice. Red line indicates 5bp deletion in 129S1/SvImJ mice. (G) Comparison of the length of intron sequences obtained from *Casp11* 5bp deletion PCR product in C57BL/6J, CBA/J, and 129S1/SvImJ mice. Data are representative of 3-5 mice per condition. Data were analyzed by unpaired Student’s t-test and are shown as average ± SEM; *p ≤ 0.05, **p ≤ 0.01, ***p ≤ 0.001.

**Figure S4 (related to Figure 4).**

(A) Representative confocal immunofluorescence image of the ileum submucosal layer of *Cx3cr1*^GFP^ mice treated with anti-CSF1R or IgG isotype control antibody by intraperitoneal injections. Scale bar = 50 µm. Images representative of at least n=3 mice per condition. (B) Representative confocal immunofluorescence image of the ileum myenteric plexus of C57BL/6J mice on day 7 post treatment with anti-CSF1R or isotype control antibody. Scale bar = 50 µm. Images representative of at least n=5 mice per condition. (C) Quantification of MM intercalation into ileum myenteric plexus ganglia as assessed by confocal immunofluorescence imaging and calculations based on Imaris software surface functions. Samples obtained at 3 hours post oral gavage of PBS or *spiB*; (D-F) CCR2^-/-^ mice were orally infected with 10^9^ CFU of *Salmonella spiB* (D) Quantification of fecal CFU on the indicated days post infection; (E) Gastrointestinal motility activity as assessed by total intestinal transit time on the indicated days post infection; experiments were ended at 450 min (dashed line); (F) Quantification of neurons of the ileum myenteric plexus as assessed by immunofluorescence staining (ANNA-1). Data are representative of at least n=4 per condition. At least 2 independent experiments were performed. Data were analyzed by unpaired Student’s t-test and are shown as average ± SEM; *p ≤ 0.05, **p ≤ 0.01, ***p ≤ 0.001.

**Figure S5 (related to Figures 5 and 6).**

(A) Gastrointestinal motility as assessed by total intestinal transit time of naïve LysM^Δ*Adrb2*^ and wild-type (WT) littermate control mice; experiments were ended at 450 min (dashed line); (B) Neuronal quantification as assessed by immunofluorescence staining (ANNA-1) in the ileum myenteric plexus of naïve LysM^Δ*Adrb2*^ and wild-type (WT) littermate control mice; (C) Ileal ring myography assessed on day 53 post infection of LysM^Δ*Adrb2*^ and wild-type (WT) littermate control mice; (D, E) Neuronal quantification as assessed by immunofluorescence staining (ANNA-1) in the myenteric plexus of ileal segments on day 7 post infection of C57BL/6 with *spiB* (D) receiving mock i.p. injections of PBS or IgG isotype control antibody or (D) following isoflurane anesthesia for 15 min 1 hour prior to infection. (D, E) Shaded area bounded by dashed lines indicates mean day 7 iEAN numbers +/- SEM of all control C57BL6/J mice analyzed in Figure 1F; (F) Neuronal quantification as assessed by immunofluorescence staining (ANNA-1) in the ileum myenteric plexus of naïve LysM^Δ*Adrb2*^ and wild-type (WT) littermate control mice. Data are representative of at least 3 mice per condition. At least 2 independent experiments were performed. Data were analyzed by unpaired Student’s t-test and are shown as average ± SEM; n.s. – not significant, *p ≤ 0.05, **p ≤ 0.01, ***p ≤ 0.001, ****p≤ 0.0001.

**Figure S6 (related to Figures 3, 4 and 6).**

(A-L) Quantification of fecal CFU on the indicated days post infection of the experiment displayed in (A) Figure 3E; (B) Figure 4C; (C) Figure 4E; (D) Figure 6B; (E) Figure 6C; (F) Figure 6D.

## Experimental Procedures

### Mice

C57BL/6J (000664), *Casp1*^-/-^ *Casp11*^-/-^(B6N.129S2-Casp1tm1Flv/J, 016621), CCR2^-/-^ (B6.129S4-*Ccr2^tm1lfc^*/J, 004999), LysM^Cre^ (B6.129P2-*Lyz2^tm1(cre)Ifo^*/J), *Arg1*^flox/flox^ (C57BL/6-*Arg1^tm1Pmu^*/J), Rosa26^tdTomato^ (B6.Cg-Gt(ROSA)26Sor^tm14(CAG-tdTomato)Hze^/J), VGLUT2^Cre^ (*Slc17a6^tm2(cre)Lowl^*/J), *Rpl22*^HA^ (B6N.129-Rpl22^tm1.1Psam^/J), 129S1(129S1/SvImJ), CBA/J, *Cx3cr1*^GFP^(*Cx3cr1*^tm1Litt/LittJ^), *Snap25*^Cre^(*Snap25^tm2.1(cre)Hze^*) were purchased from The Jackson Laboratories and maintained in our facilities. Nestin^GFP^ (Tg(Nes-EGFP)33Enik, were generously provided by P. Frenette and G. Enikolopov, and *Adrb2*^flox/flox^ (Adrb2^tm1Kry^) by G. Karsenty. Mouse lines were interbred in our facilities to obtain the final strains described in the text. Genotyping was performed according to the protocols established for the respective strains by The Jackson Laboratories, or personal communication with the donating investigators. Mice were maintained at the Rockefeller University animal facilities under specific pathogen-free conditions. Mice were fed a standard chow diet and used at 7–12 weeks of age for most experiments. Animal care and experimentation were consistent with NIH guidelines and were approved by the Institutional Animal Care and Use Committee at the Rockefeller University.

### Microorganisms

*Salmonella enterica* serovar Typhimurium (SL1344) and its mutant *spiB* were used for infection experiments and cultured prior to infection as described below. *Yersinia pseudotuberculosis* (IP32777) was cultured prior to infection as described below. *Toxoplasma gondii* was maintained in our lab by periodically infecting C57BL/6 mice with 5 cysts administrated intraperitoneally (i.p.). Cysts were isolated from brain tissue 30 days after infection. *Trypanosoma cruzi*. The *T. cruzi* Y strain was maintained by serial passage from mouse to mouse.

## Method Details

### Infections

*Salmonella enterica* Typhimurium. For infections with *Salmonella spiB*, mice were pre-treated with a single dose of Streptomycin (20 mg/mouse dissolved in 100 µl of DPBS) administered by oral gavage 18-24 hours prior to infection Mice were then orally inoculated with 10^9^ CFU of *spiB*. For re-infection experiments, mice were subjected to *spiB* infection as described above. 1 week post clearance of *spiB* from the feces, mice were fasted for hours and infected with 10^6^ CFU of wild-type *Salmonella enterica* Typhimurium (SL1344). For all *Salmonella* infections, a single aliquot of either strain of *Salmonella* was grown in 3 ml of LB overnight at 37°C with agitation. Bacteria were then sub-cultured (1:300) into 3 ml of LB for 3.5 hours at 37°C with agitation, and diluted to final concentration in 1 ml of DPBS. Bacteria were inoculated by gavage into recipient mice in a total volume of 100 µl. For experiments using heat-killed *spiB*, samples subjected to heat treatment (95°C for 10 min in water bath), prior to gavage and successful inactivation was confirmed by plating a serial dilution made from the suspension and then 5 µL onto Salmonella-Shigella plates.

*Yersinia pseudotuberculosis*. *Y. pseudotuberculosis* (strain IP32777) was grown as previously described with some adjustments (Fonseca et al., 2015). Briefly a single aliquot of the strain was grown in 3 ml of 2xYT media overnight at 28°C with vigorous agitation. Mice were fasted for 12 hours prior to infection with 10^8^ CFU by oral gavage.

*Toxoplasma gondii*. *T. gondii* was maintained in the lab by periodically infecting mice administered by intraperitoneal injection of 5 cysts/mouse in a total volume of 100 µl of DPBS.

*Trypanosoma cruzi* (*T. cruzi, Y strain)*. For each experiment, blood from an infected mouse was collected, parasites quantified and naïve recipient mice infected by intraperitoneal injection of 10^4^ parasites. Infected mice were sacrificed for tissue analysis at day 22 post infection, when the parasite load reaches a plateau (Arantes et al., 2004).

### Antibodies and flow cytometry

*Antibodies used for whole-mount immunofluorescence imaging.* The following primary antibodies were used, and unless otherwise indicated concentrations apply to all staining techniques: SST (1:200, Millipore Sigma, MAB354), ANNA-1 (1:200,000, Gift of Dr. Vanda A. Lennon), HA (1:400, Cell Signaling Technologies, 3724S), nNOS (1:200, ABCAM, ab76067), MHC II (1:400, Millipore Sigma, MABF33), ASC (1:200, Adipogen, AL177), GFP (1:400, Nacalai, GF090R). Fluorophore-conjugated secondary antibodies were either H&L or Fab (Thermo Fisher Scientific) at a consistent concentration of 1:400 in the following species and colors: Goat anti-rabbit (AF488/568/647), goat anti-rat (AF488/647), goat anti-chicken (AF488/568/647), goat anti-human (AF568/647), *Antibodies used for cell sorting of intestinal macrophages:* Fluorescent-dye-conjugated antibodies were purchased from BD-Pharmingen (USA) (anti-CD45.2, 104; anti-CD45R, RA3-6B2); eBioscience (USA) (anti-CD103, 2E7; anti-MHC II, M5; anti-F4/80, BM8; anti-CD11b, M1/70; anti-CD11c, N418; anti-Siglec F, E50-2440; anti-CD3e,145-2C11; anti-Ly6G, RB6-8C5) or BioLegend (USA) (anti-CD64 X54-5/7.1). Live/Dead staining was performed using Aqua fixable dead cell stain (Invitrogen). Macrophages were sorted as Aqua–CD45+Lin–(CD3–B220–Siglec F–LY6G–) MHCII+F4/80+CD11B+CD11C+CD103–) using a FACS Aria cell sorter flow cytometer (Becton Dickinson).

### Intestine dissection

Mice were sacrificed and duodenum (1 cm moving distal from the gastroduodenal junction), ileum (1 cm moving proximal from the ileocecal junction), or colon (4 cm moving proximal from the rectum) was removed. For dissection of the *muscularis*, following the above procedures, the intestinal tissue was placed on a chilled aluminum block with the serosa facing up (Gabanyi et al., 2016). Curved forceps were then used to carefully remove the *muscularis*.

### Nodose ganglion dissection

Mice were sacrificed and the ventral neck surface was cut open. Associated muscle was removed by blunt dissection to expose the trachea and the nodose ganglion was then located by following the vagus nerve along the carotid artery to the base of the skull. Fine scissors were used to cut the vagus nerve below the nodose ganglion and superior to the jugular ganglion.

### Colony forming unit counting

Fecal pellets from *Salmonella spiB-*infected mice were weighed and then disrupted in 400 µL of DPBS. Serial dilutions were made from the original suspension and then 5 µL of each dilution was plated onto Salmonella-Shigella plates. The plates were then incubated overnight and the number of black colonies were counted for the serial dilution with the clearest delineation of single units. This number was then multiplied by the dilution factor and by 80 to give the number of colony forming units (CFU) in the original suspension. CFU numbers were then divided by the original fecal pellet weight to give the number of CFU per mg of feces.

### Intestine motility measurements

For measurement of total intestinal transit time, mice were given an oral gavage of 6% carmine red (Sigma-Aldrich) dissolved in 0.5% methylcellulose (prepared in with sterile 0.9% NaCl). Total intestinal transit time was measured as the time from oral gavage it took for mice to pass a fecal pellet that contained carmine. Experiments were ended after 450 minutes.

### Ileal smooth muscle contractility

Ileal ring tension was assessed by organ bath (Radnoti) myography as previously described (Muller et al., 2014). Briefly, distal ileal rings were mounted and equilibrated for one hour and were distended to 0.5 g followed by 10 minutes of relaxation. The data was acquired using the PowerLab acquisition system and analyzed using LabChart Software (AD Instruments).

### Lipocalin ELISA

Lipocalin-2 levels in fecal samples were quantified using a Mouse Lipocalin-2 ELISA kit (R&D Systems) according to manufacturer’s instructions.

### Drug Administrations

*Salbutamol.* Salbutamol sulfate (Selleck Chemicals) was dissolved in sterile NaCl to a concentration of 56mg/ml, loaded into osmotic pumps and administered by subcutaneously implantation of the pumps at a final dose of 400µg/day for 14 days.

*Pan caspase inhibition.* zVAD-FMK (Selleck Chemicals) was dissolved in DMSO and administered at a dose of 50 µg/mouse by daily intraperitoneal injections over the course of a 7-day infection starting 1 day prior to infection.

*Polyamine administration*. Spermine (Sigma-Aldrich) was administered at a concentration of 2% in drinking water. Spermine-substituted drinking water was changed daily and fluid intake was monitored. Treatment was started 3 days prior to infection with *Salmonella spiB* and continued over the course of experiments.

*ODC1 inhibition.* Difluoromethylornithine (DFMO, gift from P. Woster, MSSM) was dissolved in tap water, filter-sterilized, and administered at a concentration of 4% in drinking water. DFMO-substituted water was changed daily and fluid intake was monitored. Treatment was started 4 days prior to infection with *Salmonella spiB* and continued over the course of experiments.

### Isoflurane induction

A plexiglass induction chamber was placed on a heating pad set to 37°C. Isoflurane was set at 1% with an oxygen flow of 1 liter/minute. Mice were placed in the induction chamber and monitored for loss of movement, after which a period of 15 minutes was elapsed. Respiration was monitored continuously during this period. Following isoflurane treatment, mice were returned to their home cage for a period of 1 hour prior to receiving Streptomycin pre-treatment, followed by *spiB* infection as described above.

### Sham intraperitoneal injections

Sham intraperitoneal injections of DPBS or IgG (isotype control from MM depletion experiments, see below) were performed by mimicking the anti-CSF1R MM depletion protocol, by intraperitoneal injection of 200 µl of DPBS or IgG/ mouse (100 µl/flank). Injections were performed 12 hours prior to *spiB* infection, and an additional injection was performed at day 3.5 post infection. Mice were sacrificed and tissue was analyzed at 7 dpi.

### Subcutaneous pump implantation

Mice were anesthetized using isoflurane and, under sterile conditions, a small incision was made on the dorsal side of the neck. An Alzet pump (Model 1002) filled with Salbutamol sulfate prepared as described above was placed under the skin and moved back towards the right flank. The incision was then closed with surgical wound clips (Kent Scientific). Mice were then left to rest for 7 days before infection with *spiB*.

### Anti-CSF1R (ASF98) antibody production

Anti-CSF1R was produced by suspension culture of the ASF98 hybridoma cell line (gift from Miriam Merad). Cells were thawed and passaged twice (1:10) in PFHM-II (Thermo Fisher) media supplemented with P/S (Thermo Fisher). Cells were then seeded at 5×10^6^ in 15 mL of PFHM-II in the cellular compartment of a 1000 mL bioreactor (Wheaton) and allowed to grow for 7-10 days. The cell compartment was then harvested, spun down and the supernatant collected. Supernatant was filtered and antibody was purified from the supernatant by affinity purification using protein G sepharose (GE Healthcare) in a gravity column (Bio-Rad). Briefly, the protein G was equilibrated with 100 mL binding buffer (Thermo Fisher Scientific) and supernatant was loaded onto and run through the column. The column was then washed with 200 mL of binding buffer and eluted with 10 mL elution buffer (Thermo Fisher Scientific). Antibody was eluted in fractions directly into 100 µl 1 M Tris-HCL pH 8 (Invitrogen) and the concentrations were measured on a Nanodrop 2000 spectrophotometer (Thermo Fisher Scientific). Fractions were combined and dialyzed in 2 L DPBS for 2 days at 4°C using 14,000 MWCO dialysis tubing (Spectrum Labs) changing once. The antibody was then concentrated using centrifugal filters (30,000 MW, Amicon) and stored at 4°C until use.

### MM depletion using anti-CSF1R

Anti-CSF1R was diluted in sterile DPBS to a final concentration of 6.25 mg/ml and 50 mg/kg of anti-CSF1R or isotype control (IgG from rat serum, Sigma-Aldrich) were administered by i.p. injections. Mice were previously habituated to i.p. injections for at least 5 days prior to the start of depletion and *spiB* infection. For all experiments, depletion was performed twice over the course of 7 days (day 0 and day 3.5). For infection experiments, depletion was performed 12 hours prior to infection. Animals were then subjected to the *Salmonella spiB* infection protocol as described above, and received and additional dose of either anti-CSF1R or isotype control 3.5 days after the initial dose. Animals were sacrificed at 7 dpi and intestinal tissue dissected for analysis.

### Immunocytochemistry

Cuprolinic blue staining for visualization of enteric neurons was performed as previously described (Holst and Powley, 1995). The method was utilized for quantification of enteric neurons in *T. gondii* and *T. cruzi* infection experiments (Figure S1).

### Whole-mount intestine immunofluorescence

Briefly, mice were sacrificed by cervical dislocation and the small intestine was removed and placed in HBSS Mg^2+^Ca^2+^(Gibco) + 5% FCS. The intestine was cut open longitudinally and the luminal contents washed away in DPBS. The *muscularis* was then carefully dissected away from the underlying mucosa in one intact sheet. The tissue was pinned down in a plate coated with Sylgard and then fixed for O/N with 4% PFA with gentle agitation. After washing in DPBS whole mount samples were then permeabilized first in 0.5% Triton X-100/0.05% Tween-20/4µg heparin (PTxwH) for 2 hours at room temperature (RT) with gentle shaking. Samples were then blocked for 2 hours in blocking buffer (PTxwH with 5% bovine serum albumin/5% donkey/goat serum) for for 2 hours at RT with gentle agitation. Antibodies were added to the blocking buffer at appropriate concentrations and incubated for 2 days at 4°C. After primary incubation the tissue was washed 4 times in PTxwH and then incubated in blocking buffer with secondary antibody at concentrations within the primary antibody range for 2 hours at RT. Samples were again washed washed 4 times in PTxwH and then mounted with FluoroMount G on slides with 1 ½ coverslips. Slides were kept in the dark at 4°C until they were imaged.

### Confocal imaging

Whole mount intestine samples were imaged on an inverted LSM 880 NLO laser scanning confocal and multiphoton microscope (Zeiss) and on an inverted TCS SP8 laser scanning confocal microscope (Leica).

### Neuronal quantification

A minimum of 10 images were randomly acquired across a piece of whole mount *muscularis*. These images were then opened in ImageJ, and the cell counter was used to count the number of ANNA-1+ cells in a given field. This number was then multiplied by a factor of 2.95 (20x objective) or 3.125 (25x objective), to calculate the number of counted neurons per square millimeter (mm^2^). The average of 10 (or more) images were then calculated and plotted. Thus, every point on a given graph corresponds to a single animal. For subtypes, the number of nNOS, SST, and tdTomato+ neurons were also counted. These numbers were then reported as both number per mm^2^ and percent of ANNA-1+ neurons.

### RiboTag

Heterozygous or homozygous *Snap25*^RPL22HA^ were used for TRAP-seq analysis as no differences were found between either genotype. For intestine immunoprecipitation (IP) mice were sacrificed and tissue remove and divided as above. Samples were washed of fecal contents in PBS with cycloheximide (0.2 mg/mL) (PBS/CHX). Mesenteric fat was removed and the *muscularis* was separated from the mucosa as described above and samples were washed 5 times in PBS/CHX. For nodose tissues were isolated as described above. The RiboTag immunoprecipitation protocol (http://depts.washington.edu/mcklab/RiboTagIPprotocol2014.pdf) was then followed with the following modifications: All samples were homogenized by hand with a dounce homogenizer in 2.5 mL supplemented homogenization buffer (changes per 2.5 mL: 50 µl Protease Inhibitor, 75 µl heparin (100 mg/mL stock), 25 µl SUPERase In RNase Inhibitor). Samples were then centrifuged for 10 minutes at 10,000 G, after which 800 µl of supernatant was removed and 5µL of anti-HA antibody (Abcam, ab9110) was added. Samples were kept rotating at 4°C with antibody for 1 hour. 200 µl of Thermo Protein magnetic A/G beads were washed with homogenization buffer, added to the sample, and kept rotating for 30 minutes at 4°C. The beads were washed four times with high-salt buffer and samples were eluted with 100 µl of PicoPure lysis buffer. RNA was extracted using the Arcturus PicoPure RNA isolation kit (Applied Biosystems) according to the manufacturer’s instructions.

### RNA-sequencing

RNA libraries were prepared using SMARTer Ultra Low Input RNA (ClonTech Labs) and sequenced using 75 base-pair single end reads on a NextSeq 500 instrument (Illumina). Reads were aligned using Kallisto (Bray et al., 2016) to Mouse Ensembl v91. Transcript abundance files were then used in the DESeq2 R package, which was used for all downstream differential expression analysis and generation of volcano plots. Differentially expressed genes between samples were compared with a cutoff of log_2_ Fold Change > 1.

### Cryosectioning of fresh-frozen tissue

Mice were given a lethal dose of isoflurane and then perfused with 30 ml ice cold DPBS. The ileum and colon were removed and flushed of luminal contents. The tissue was then transferred to OCT, positioned, flash frozen, moved to −20 for 2 hours, and finally moved to −80 for long term storage. Prior to cryosectioning, OCT blocks were equilibrated in the cryostat for 20 minutes. Sections were cut at 15 µm, put onto FrostPlus slides, and left to dry for 10-15 minutes. Slides were then transferred to dry ice and finally to −80°C before RNAscope® processing.

### RNA fluorescence *in situ* hybridization (FISH) using RNAscope® technology

RNAscope® *in situ* hybridization was performed using probes against *N*lrp6 and *Elavl4* on 15 µm sections of fresh frozen ileum or colon tissue isolated from C57BL/6J mice according to the manufacturer’s instruction with the following modification, tissue was pretreated with Protease IV for 20 min at RT. Samples were then mounted in Fluoromount-G with DAPI and 1 ½ coverslips were applied.

### Macrophage intercalation calculation

Raw data as imported from the microscope was used for all cell identification and subsequent analyses. Some post – analysis pseudo-color adjustment was performed for individual images to account for differences in auto-fluorescence and labeling. *Imaris* (Bitplane AG) software was used for cell identification, using the “*Surfaces*” algorithm as described by the manufacturer. A surface was created based on anti-ANNA-1 (neuron) staining to define neuronal ganglia. A masked channel was then created to capture fluorescence in other channels within the neuronal ganglia volume. A second surface was created based on anti-MHCII (macrophage) staining only within the masked channel. Volume statistics on both neuron (total) and macrophage (neuronal ganglia masked) surfaces were then exported. Percent macrophage intercalation per image was calculated as:

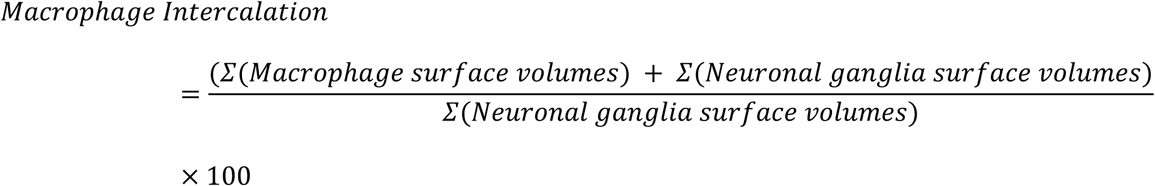

The average of at least 10 random macrophage intercalation percentages was used to determine the percentage for each individual animal.

### 3D image reconstruction for MM ganglionic intercalation calculations

Images were adjusted post hoc using Imaris x64 software (version 9.1 Bitplane) and 3D reconstructions were recorded as mp4 video files. Optical slices were taken using the orthoslicer or oblique slicer tools.

### Determination of caspase 11 mutation status by PCR and Sanger Sequencing

DNA was extracted from ear pieces of mice using QuickExtract™ DNA Extraction Solution (Lucigen). Target sequences were amplified using the following primers: Forward 5’-AGGCATATCTATAATCCCTTCACTG-3’; Reverse 5’-GAATATATCAAAGAGATGACAAGAGC-3’. The following PCR conditions were used: 4 min 94°C, 1 min 94°C, 0.5 min 58°C, 1 min 72°C, 7 min 72°C, and end at 12°C. Samples were loaded on 3% agarose gels, and bands from all strains were compared to samples from C57BL/6J and 129S mice. The 5 bp deletion in *Casp11* described in the 129S1 strain fragment runs slightly higher than the wild-type fragment at a height of approximately 220 bp (Vanden Berghe et al., 2015). *Casp11* mutation status was further confirmed in C57BL/6J, CBA/J and 129S1 mice by Sanger sequencing of a gel-purified PCR product using primers 5′-CAGTATTATTATTGGTGATGCAAATG-3′ and 5′-GGAATATATCAAAGAGATGACAAGAGC-3′. *Casp11* mutation status was also confirmed in further relevant mouse strains utilized in this study.

### Single Cell Suspension of Intestinal Macrophages

Mice were euthanized, and the small intestine was carefully removed, cleaned, cut open longitudinally and washed 2X in HBSS Mg^2+^Ca^2^Gibco) and 1X in HBSS Mg^2+^Ca^2^ with 1mM DTT (Sigma-Aldrich). The tissue was cut in two and the *muscularis* region was carefully dissected from the underlying mucosa. *Muscularis* tissue was then finely cut and digested in HBSS Mg^2+^Ca^2+^+ 5% FBS + 1x NaPyr + 25mM HEPES

+ 50 μg/ml DNase I (Roche) + 400U/ml Collagenase D (Roche) + 2.5U/ml Dispase (Corning) at 37°C. The *muscularis* was digested for 40 min. The tissue was then homogenized with an 18-gauge needle, filtered through a 70 μm cell strainer and washed with HBSS Mg^2+^Ca^2^. Cells were incubated with Fc block and antibodies against the indicated cell surface markers in FACS buffer (PBS, 1% BSA, 10 mM EDTA, 0.02% NaN_3_).

### Quantitative PCR

Total RNA was isolated using TRIzol^TM^ (Invitrogen), from which cDNA libraries were reverse transcribed using Superscript II (Invitrogen) and random primers following the instructions provided by the manufacturer. Quantitative PCR was performed using SYBR green (Bio-Rad Laboratories). Data were collected and analyzed on a QuantStudio 3 (Thermo Scientific). Th *Rpl32* housekeeping gene was used to normalize samples. The following primers were used: Rpl32 forward 5’-ACAATGTCAAGGAGCTGGAG-3’, Rpl32 reverse 5’-TTGGGATTGGTGACTCTGATG-3’, *Arg1* forward 5’-CTCCAAGCCAAAGTCCTTAGAG-3’, *Arg1* reverse 5’-AGGAGCTGTCATTAGGGACATC-3’, *Ym1* forward 5’-AGACTTGCGTGACTATGAAGCATT-3’, *Ym1* reverse 5’-GCAGGTCCAAACTTCCATCCTC-3’

### Statistical analysis

Significance levels indicated are as follows: * *P* < 0.05, ** *P* < 0.01, *** *P* < 0.001, **** *P* < 0.0001. All data are presented as mean ± SD or mean ± SEM. All statistical tests used were two-tailed. The experiments were not randomized and no statistical methods were used to predetermine sample size. Multivariate data was analyzed by one-way ANOVA and Tukey’s multiple comparisons post hoc test. Comparisons between two conditions were analyzed by unpaired Student’s t-test. We used GraphPad PRISM version 8.0 and R 3.4.3 for generation of graphs and statistics.

### Data and Software Availability

Data generated by RNA sequencing will be deposited in the NIH GEO database upon acceptance for publication. All software used is available online, either freely or from a commercial supplier and is summarized in the Key Resources table. Algorithms described above are purely mathematical calculations and can be performed using any software. No new software was written for this project.

