## Supplementary figures and images for "Adrenergic signaling in muscularis macrophages limits neuronal death following enteric infection"

### Supplemental Figure 1

#### Figure S1 (related to Figure 1)

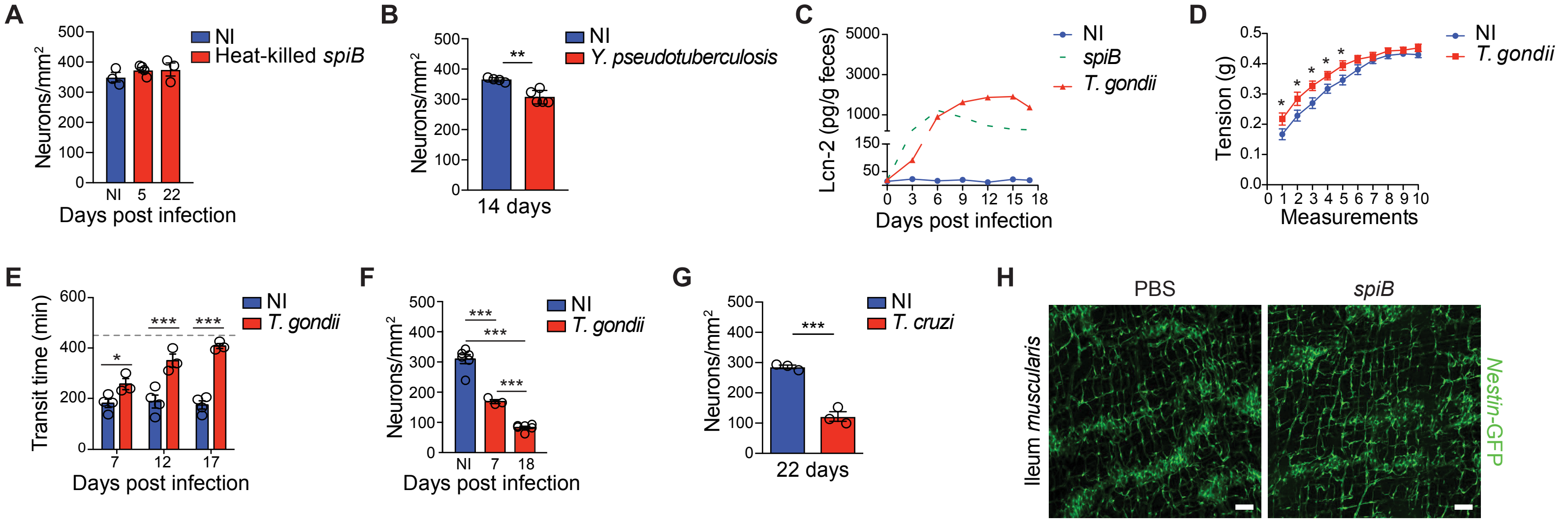

### Supplemental Figure 2

**Figure S2 (related to Figure 2).**

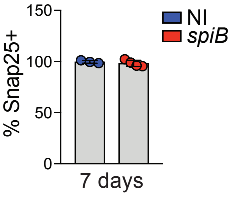

### Supplemental Figure 4

#### Figure S4 (related to Figure 4)

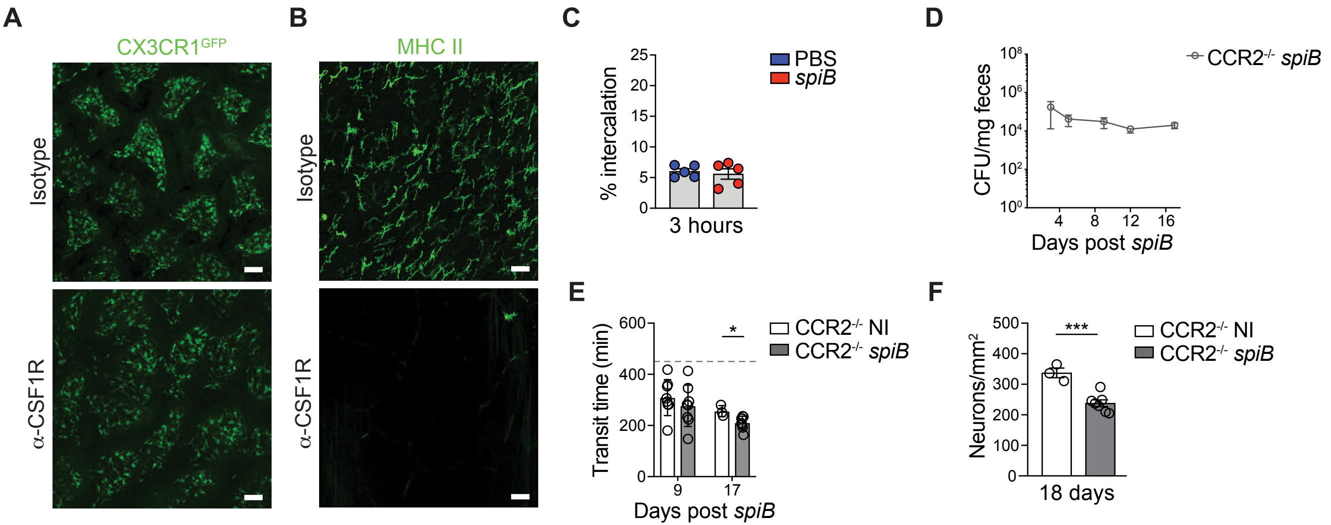

### Supplemental Figure 5

#### Figure S5 (related to Figures 5 and 6)

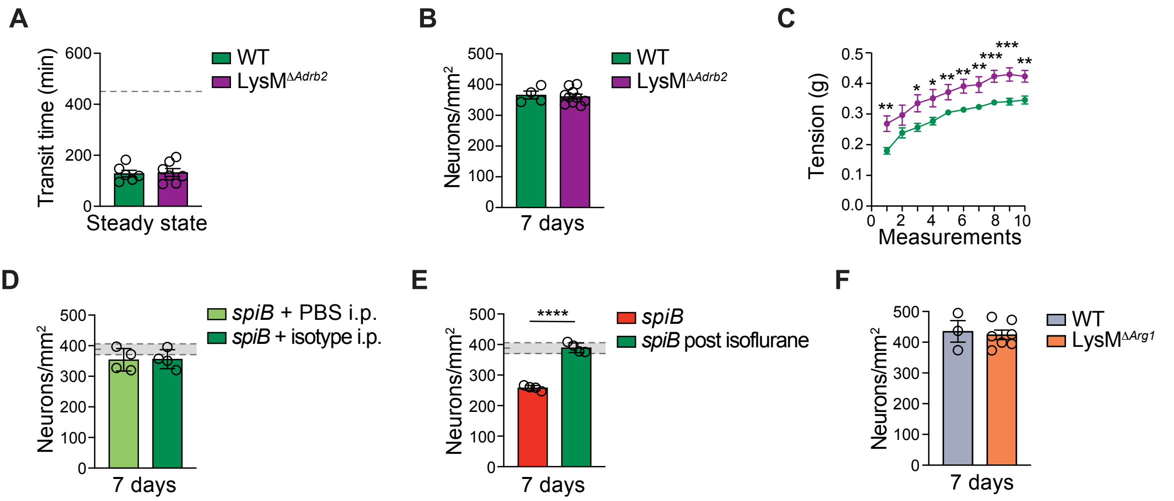

### Supplemental Figure 6

#### Figure S6 (related to Figures 3, 4 and 6).

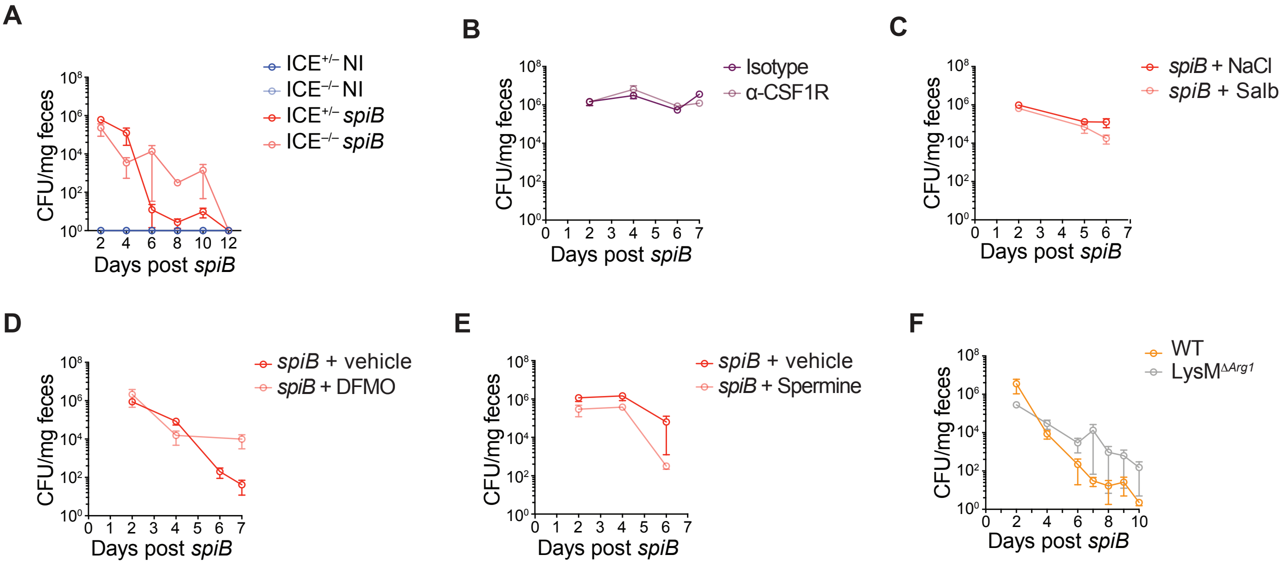
