## Supplemental Figure 3 for "Adrenergic signaling in muscularis macrophages limits neuronal death following enteric infection"

### Figure S3 (related to Figure 3)

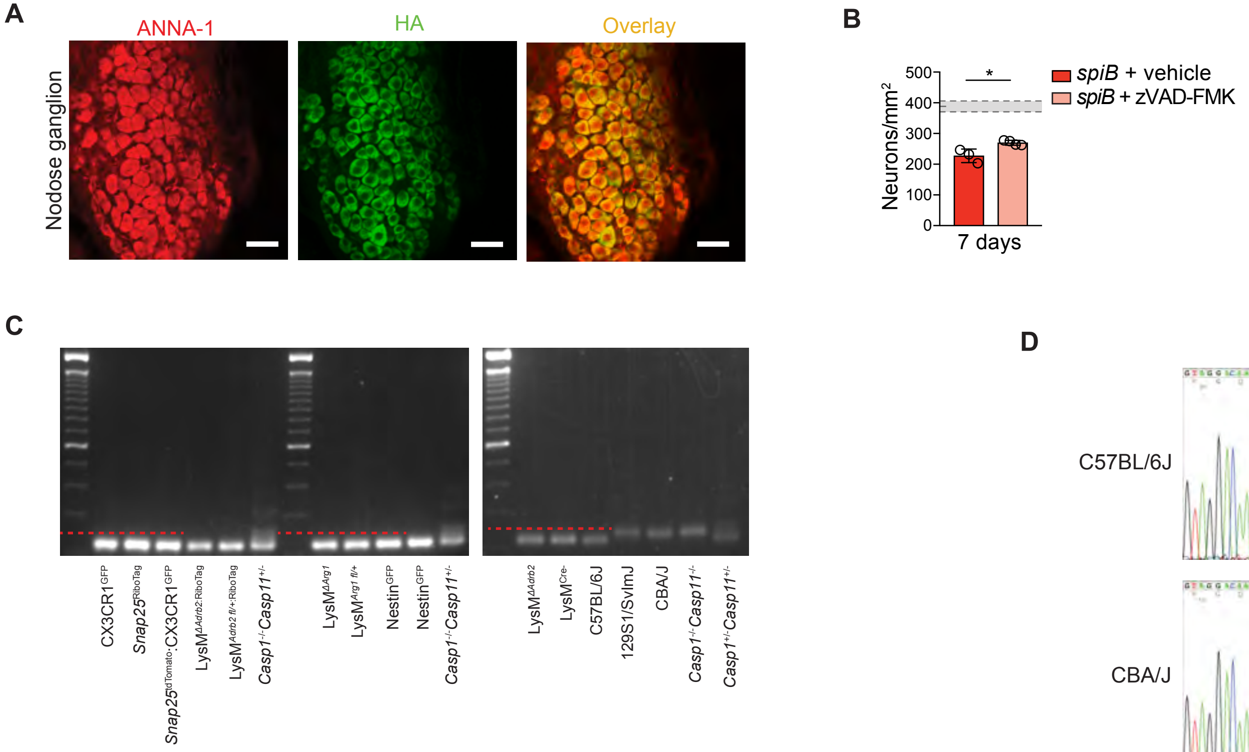


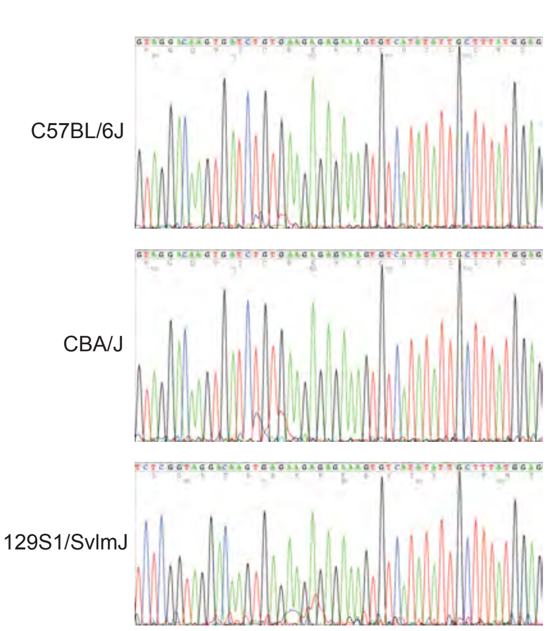


**E**

| | |
|---|---|
| C57BL/6J | GTAGGACAAGTGATCTGTGAAGAGAGAAAGTGTCATATATTGCTTTATGGAG |
| CBA/J | GTAGGACAAGTGATCTGTGAAGAGAGAAAGTGTCATATATTGCTTTATGGAG |
| 129S1/SvlmJ | GTAGGACAAGTGA —— GAAGAGAGAAAGTGTCATATATTGCTTTATGGAG |

**F**

| | |
|---|---|
| C57BL/6J(83bp) | TTTTTTTTTTTTTTTTTTTTTTTTTTTTTTTTTTTTTAAAAAACCCGGGGCCCCGGGGGGGGGGGGTTTAAAAATTCCAAAAAN |
| CBA/J (98bp) | TTTTTTTTTTTTTTTTTTTTTTTTTTTTTTTTTTTTTTTTTTTTTTTTTTTTTTTAAACCCCCCGGGCCCCCGGGGGGGGGGGGTTTATAAAAAAAAAAAAAN |
| 129S1/SvlmJ(107bp) | TTTTTTTTTGTTTTTTTTTTTTTTTTTTTTTTTTTTTTTTTTTTTTTTTTTTTTTTTTTTTTTACCCCCCCCGGGGCCCCGGGGGGGGGGGGTTTTTTAAATAAAAAAAAN |
